# Opportunistic bacteria of grapevine crown galls are equipped with the genomic repertoire for opine utilization

**DOI:** 10.1101/2023.06.20.545721

**Authors:** Hanna Faist, Markus J. Ankenbrand, Wiebke Sickel, Ute Hentschel, Alexander Keller, Rosalia Deeken

## Abstract

Young grapevines (*Vitis vinifera*) suffer and eventually can die from the crown gall (CG) disease caused by the plant pathogen *Allorhizobium vitis (Rhizobiaceae)*. Virulent members of *A. vitis* harbour a tumor-inducing (Ti) plasmid and induce formation of crown galls (CGs) due to the oncogenes encoded on the transfer-DNA (T-DNA). Expression of oncogenes in transformed host cells induce unregulated cell proliferation, metabolic and physiological changes. The CG produces opines uncommon to plants, which provide an important nutrient source for *A. vitis* harbouring opine catabolism enzymes. CGs host a distinct bacterial community and the mechanisms establishing a CG-specific bacterial community are currently unknown. Thus, we were interested in whether genes homologous to those of the Ti-plasmid coexist in the genomes of the microbial species coexisting in CGs.

We isolated eight bacterial strains from grapevine CGs, sequenced their genomes and tested their virulence and opine utilization ability in bioassays. In addition, the eight genome sequences were compared to 34 published bacterial genomes, including closely related plant associated bacteria not from CGs. Homologous genes for virulence and opine anabolism were only present in the virulent Rhizobiaceae. By contrast, homologs of the opine catabolism genes were present in all strains including the non-virulent members of the Rhizobiaceae and non-Rhizobiaceae. Gene neighbourhood and sequence identity of the opine degradation cluster of virulent and non-virulent strains together with the results of the opine utilization assay support the important role of opine utilization for co-colonization in CGs, thereby shaping the CG community.

**Significance statement:** Virulent *Allorhizobium vitis* causes crown galls on grapevines which reduce plant vigour, yield, and cannot be cured. Non-virulent agrobacteria have been used as biocontrol agents to reduce the virulence potential within a crown gall and disease symptoms. We wanted to know if and how in nature this biocontrol concept is accomplished. We found virulent *Allorhizobium* along with non-virulent *Agrobacterium*, or *Pseudomonas* in the same tumours. Both harboured the catabolism genes in their genomes and metabolized the *quorum sensing* molecule opine. Thus, in nature it seems common that virulent and non-virulent species coexist in a crown gall and that the avirulent members control the virulence potential of the crown gall community by reducing the opine levels.

## Introduction

*Allorhizobium vitis* (*Rhizobiaceae*), former *Agrobacterium vitis* or *Agrobacterium tumefaciens* biovar 3, is the causal pathogen of grapevine crown galls (CGs) which hamper plant growth and yield (Ferreira et al. 1992; Schroth et al. 1988). Overall, the family *Rhizobiaceae* contains both virulent and non-virulent species (Bien et al. 1990; Chandrasekaran et al. 2019) and *Rhizobiaceae* are associated with different grapevine tissues (Burr and Katz 1983). Virulence is encoded by the bacterial tumor-inducing plasmid (Ti-plasmid; Wikipedia contributors 2023) that consists of the transfer DNA (T-DNA), the virulence gene operon (including *vir*-genes), genes encoding opine utilization enzymes as well as a bacterial backbone region that regulates replication and conjugation of the Ti-Plasmid (Chen and Xie 2011; Gordon and Christie 2014). The Vir-proteins guide the transfer of the T-DNA into the plant nucleus and enable integration into the plant genome. The transformed plant cells express T-DNA encoded genes, consequently leading to the production of plant growth hormones and opines (Gelvin 2010). Uncontrolled production of the plant hormones auxin and cytokinin cause plant cell proliferation and thereby tumorous growth, also referred to as crown galls (CGs; Gohlke and Deeken 2014; Escobar et al. 2001; Klee et al. 1984). CGs do not only offer space for virulent *Rhizobiaceae*, but also for a specific bacterial community which is distinct and differs from the community of a normal wound callus of the graft union (Faist et al. 2016). Moreover, a recent study on the bacterial composition of 73 CGs has shown that at least three non-*A. vitis* groups co-exist in CGs (Gan et al. 2019)

Opines produced by CG cells are a source of nitrogen and carbon for *A. vitis* and the capacity to utilize opines provides a fitness advantage over non-opine utilizing bacteria (Lang et al. 2017). For virulent *Rhizobiaceae*, opines represent not only a nutrient source, but also are involved in *quorum sensing* that regulate e.g., Ti plasmid conjugation and its distribution between bacteria (Ellis et al. 1982; Wetzel et al. 2014). Different virulent *Rhizobiaceae* transfer different opine biosynthesis genes into the plant genome. Various opines are known (Dessaux et al. 1998; Moore et al. 1997; Chilton et al. 2001) and in CGs nopaline, octopine/cucumopine, and/or vitopine/heliopine have been found of which the latter opine-type exclusively occurs in grapevine CGs (Szegedi et al. 1988; Szegedi 2003). It has been postulated that the vitopine/heliopine-type pTi’s of *A. vitis* represent a distinct group of Ti-plasmids (Szegedi et al. 1996). Indeed, the bacterial catabolism genes on the Ti-plasmid correspond to these opine-types. Nevertheless, as opines are produced by T-DNA transformed plant cells, they are a public good in a CG (Platt et al. 2012). Consequently, other bacteria of the CG community may also utilize opines as a nutrient source, promoting their enrichment in CGs. For example, some *Pseudomonas* strains isolated from CGs can utilize opines (Bergeron et al. 1990; Moore et al. 1997).

In our study, we provide the draft genome sequences of three virulent *A. vitis* isolates and five non-virulent bacterial isolates of grapevine CG communities (three *Rhizobiaceae*, one *Pseudomonas* and one *Rahnella*) and analysed these strains for virulence. In octopine and nopaline utilization bioassays we tested growth of the eight grapevine CG isolates. In addition, 34 published genomes of plant associated bacteria were included in our analyses to identify orthologous genes of the Ti-Plasmids. In our genome analysis, we focussed on the distribution of genes involved in virulence, Ti-plasmid conjugation (quorum sensing), and opine metabolism in the CG bacterial communities.

## Results

### Physical characteristics of the eight genomes

The eight *de novo* sequenced bacterial genomes belonged to isolates from five different grapevine CGs harvested in the region Franconia, Bavaria, Germany (supplementary table S1). According to EZBioCloud (Yoon, et al. 2017) the isolates CG1-CG6 belonged to the *Rhizobiaceae* family, CG7 was identified as *Pseudomonas sp*., and CG8 as *Rahnella sp.* (supplementary table S1). The screening for amplicon sequencing variants (ASVs) of the dataset published by Faist et al. 2016 revealed that the V4 regions of the 16S rRNA sequences from the isolates CG1-CG3 and CG7-CG8 were significantly enriched in CGs compared to healthy graft unions (table 1) whereasASVs of CG4-CG6 showed no significant enrichment (Faist et al. 2016).

**Table 1.**
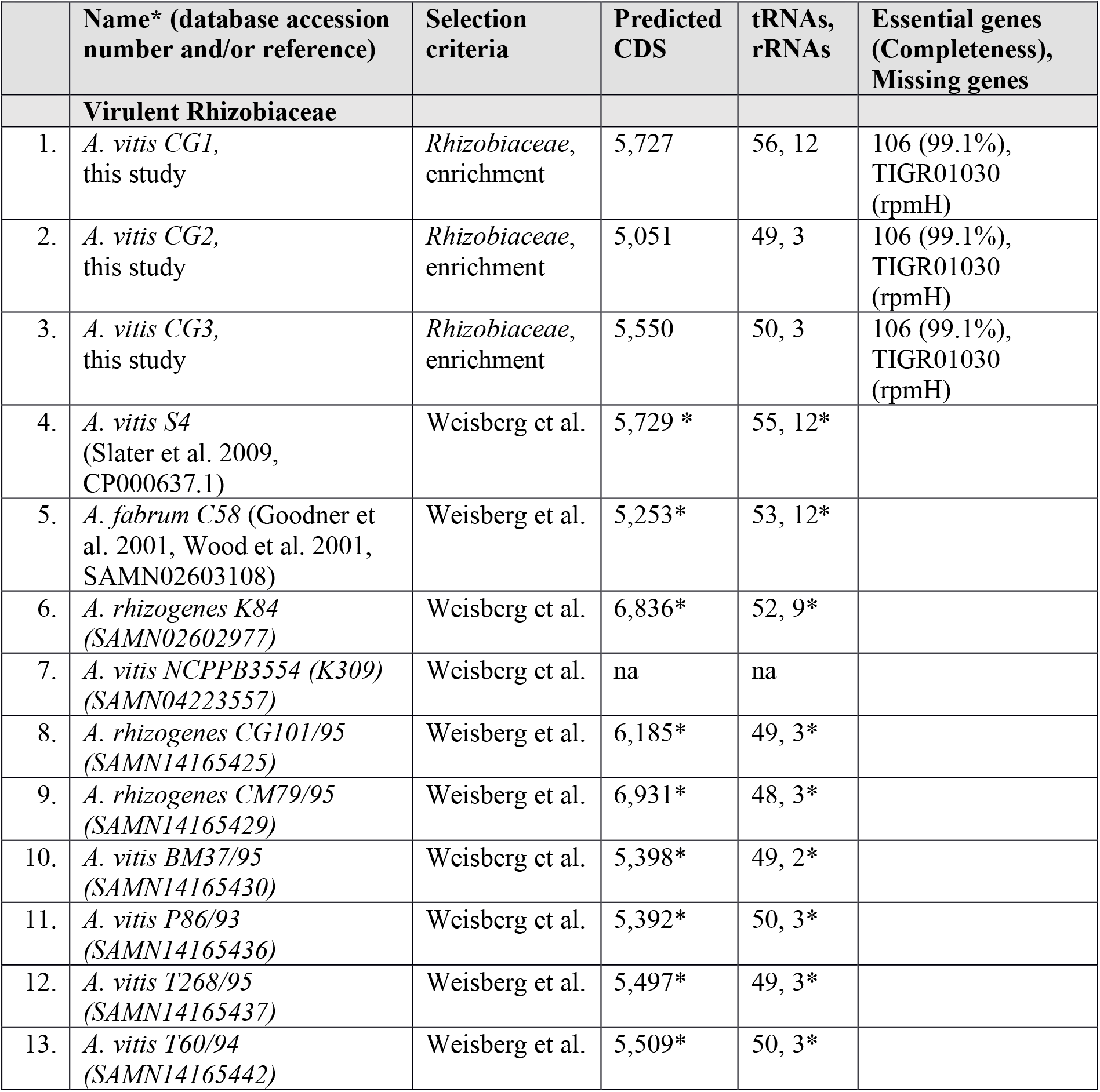
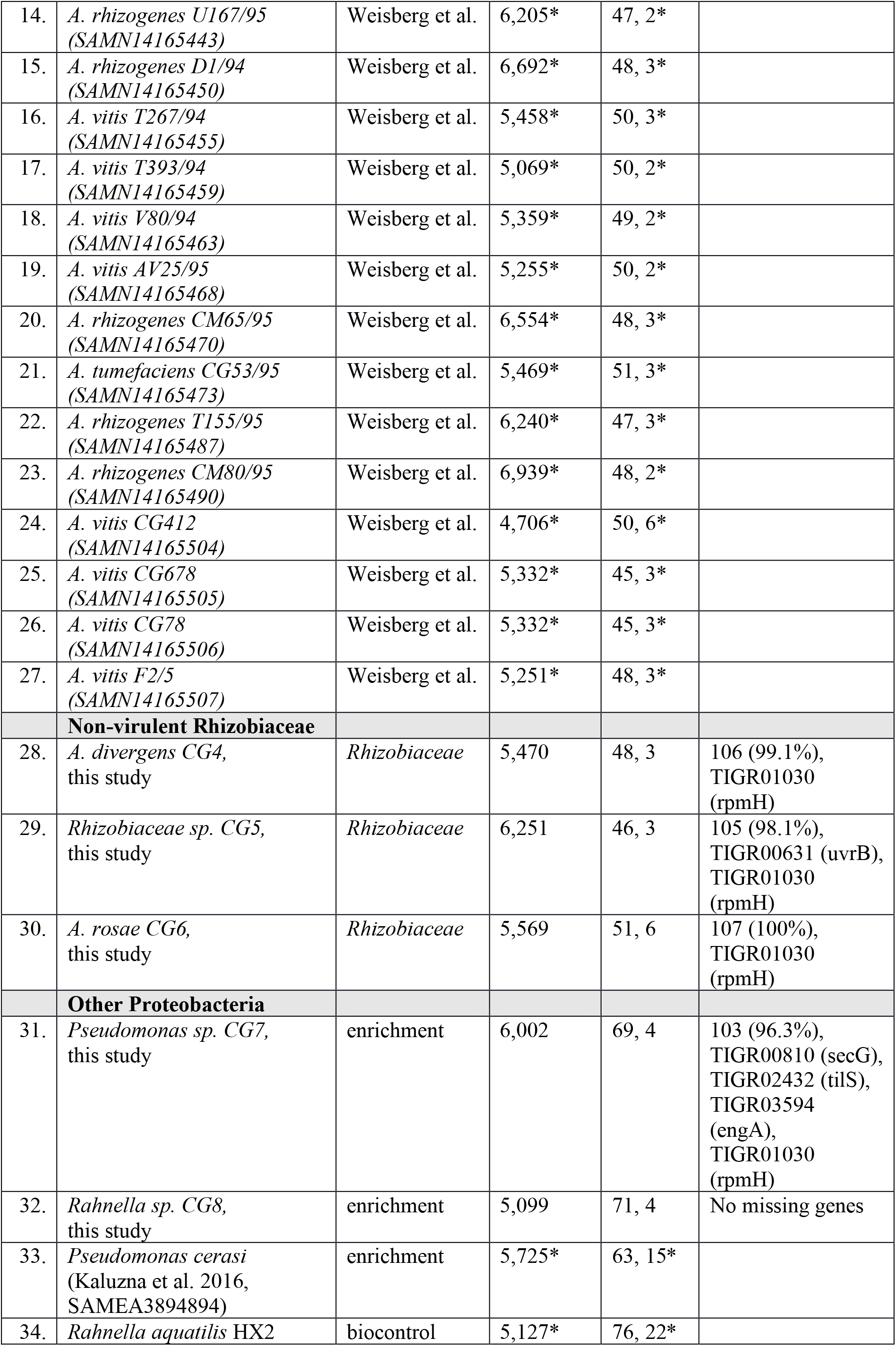
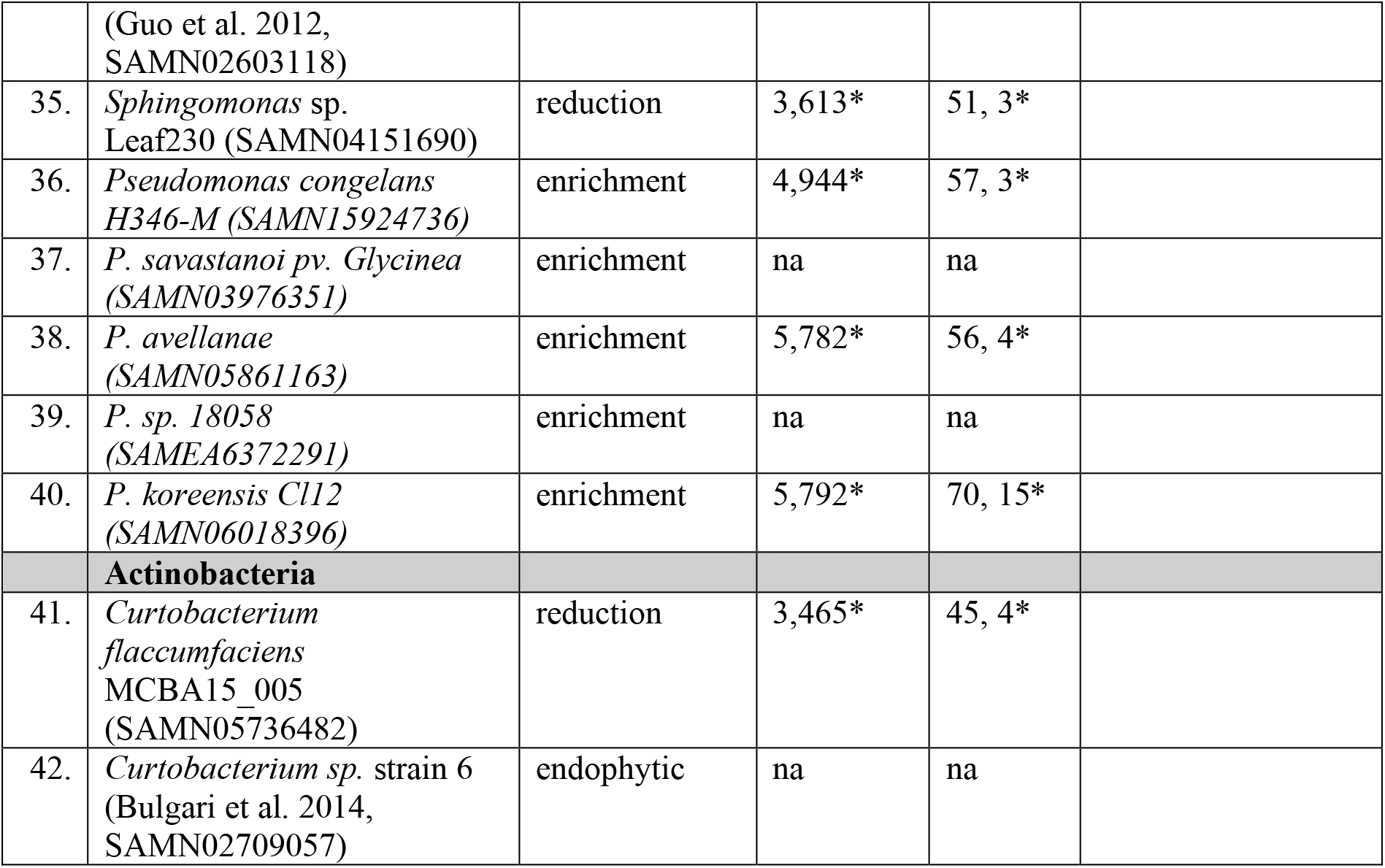
Information about the bacterial genomes analysed in this study (1.-3., 7.-11.) and the known reference sequences (4.-6., 12.-16.) used for comparison. The selection criteria for the sequences were: i) ‘enrichment’ or ‘reduction’ of the 16S rRNA V4 amplicon sequences in crown galls in springtime (Faist et al., 2016), ii) member of the *Rhizobiaceae* family, (iii) part of the agrobacterial virulence plasmid study Weisberg et al., 2020, iv) endophytic in grapevine graft unions without crown gall disease, and v) biocontrol for crown gall disease. Annotation statistics for genomes from this study or according to NCBI assembly report (genome annotation data, if available, indicated with *), CDS, coding sequences

The general features of the assembled eight draft genomes are summarized in supplementary table S2. The length of the smallest contigs, which accounted for 90% of the genome (N90 index), ranged from 17.6 kb (CG7, *Pseudomonas*) to 204 kb (CG6, *Rhizobiaceae sp.*). The *A. vitis* draft genomes (CG1-CG3) shared a GC content of around 57%, while it varied between the other *Rhizobiaceae* isolates (CG4-CG7, 55-61.5%). *Rahnella* (CG8) possessed the lowest GC content with 52.3%. Mapping the raw reads to the draft genomes resulted in ∼98% alignment rates for CG1, CG2, CG4-CG6 and about ∼85-87% for the isolates CG3, CG7, and CG8.

Inoculation assays demonstrated that the three *A. vitis* isolates (CG1-CG3) induced CG development on stems of in vitro cultivated grapevine plantlets but not the other three Rhizobiaceae isolates (CG4-CG6; supplementary fig. 1). Like the latter three, *Pseudomonas* (CG7) and *Rahnella* (CG8) did not induce CG development even in stems of the test plants *Arabidopsis thaliana* and *Nicotiana benthamiana*. Thus, the isolates CG1-CG3 were confirmed as virulent, while CG4-CG8 were non-virulent.

**Fig. 1.**
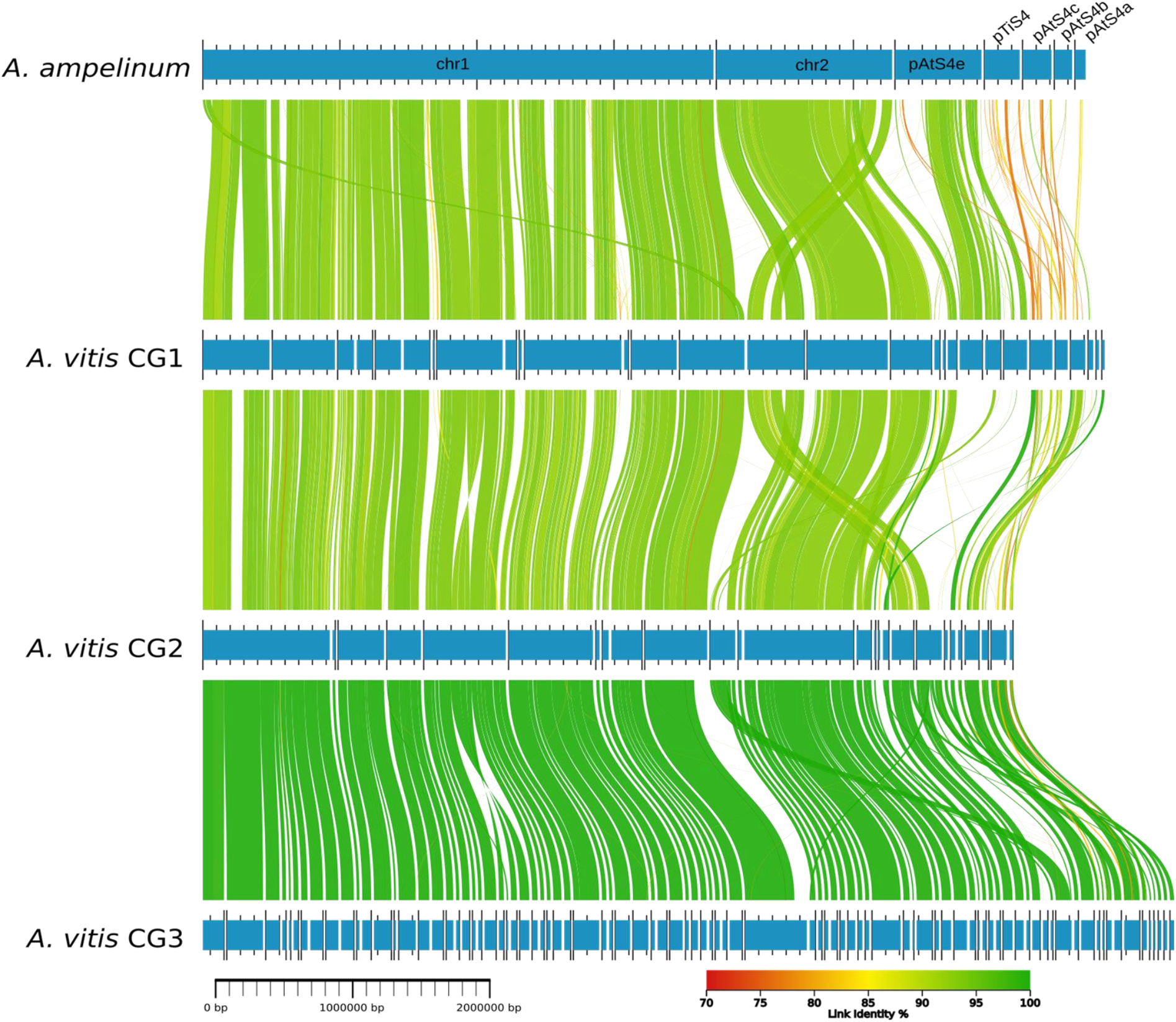
DNA alignment of *Allorhizobium vitis* genomes. Blue horizontal bars indicate the chromosomes (chr1, chr2) and plasmids (pATS4e, pTiS4, pTiS4a-c) of *A. ampelinum*, and the contigs of the *A. vitis* CG1-CG3 genome sequences. Homologous regions between genomes are connected via lines of different colors. A color gradient (70 to 100%) according to the similarity (Link identity %) is provided.

### Relationship between isolates and reference genomes

Information about the selection criteria of the *de novo* sequenced draft genomes (CG1-CG8) and the reference sequences of known plant associated bacteria used for phylogenetic relationship analysis are summarized in table 1. The numbers of predicted coding sequences (CDS) and tRNAs are similar among the isolates while the number of rRNAs varies, most likely due to the highly challenging assembly of frequently duplicated genes in draft genomes. At least 103 out of 107 essential genes were found in all *de novo* sequenced isolates indicating a completeness of the draft genomes of at least 96%.

The *Rhizobiaceae* family encompasses virulent and non-virulent members including the monophyletic groups of *Agrobacterium* and *Allorhizobium* (Gan and Savka 2018). The results of a phylogenetic tree generated on the basis of 107 housekeeping gene sequences revealed that the two non-virulent *Rhizobiaceae* isolates CG4 *(A. divergens)* as well as CG6 (*A. rosae*) belong to the clade of *Agrobacterium* while CG5 is related to *Allorhizobium* (supplementary fig. 2). The three virulentisolates CG1-CG3 formed a clade also with the genus *Allorhizobium* of which CG2-CG3 belonged to the branch of *A. vitis*. A whole genome alignment using the program AliTV performed with *A. ampelinum* and the draft genomes of the isolates CG1-GC3 revealed that the two chromosomes and the pATS4e plasmid express high homology (fig. 1, green connecting lines, >85% homology), while the pTiS4 plasmid possessed much lower homology (orange to yellow connecting lines, <85% homology). The close relationship between the virulent *A. vitis* genomes was confirmed by the number of protein families shared (fig. 2). CG1, CG2, and CG3 exclusively shared 211 protein families with *A. ampelinum* and *A. vitis* NCPPB3554 (K309) (fig. 2, arrow).

**Fig. 2.**
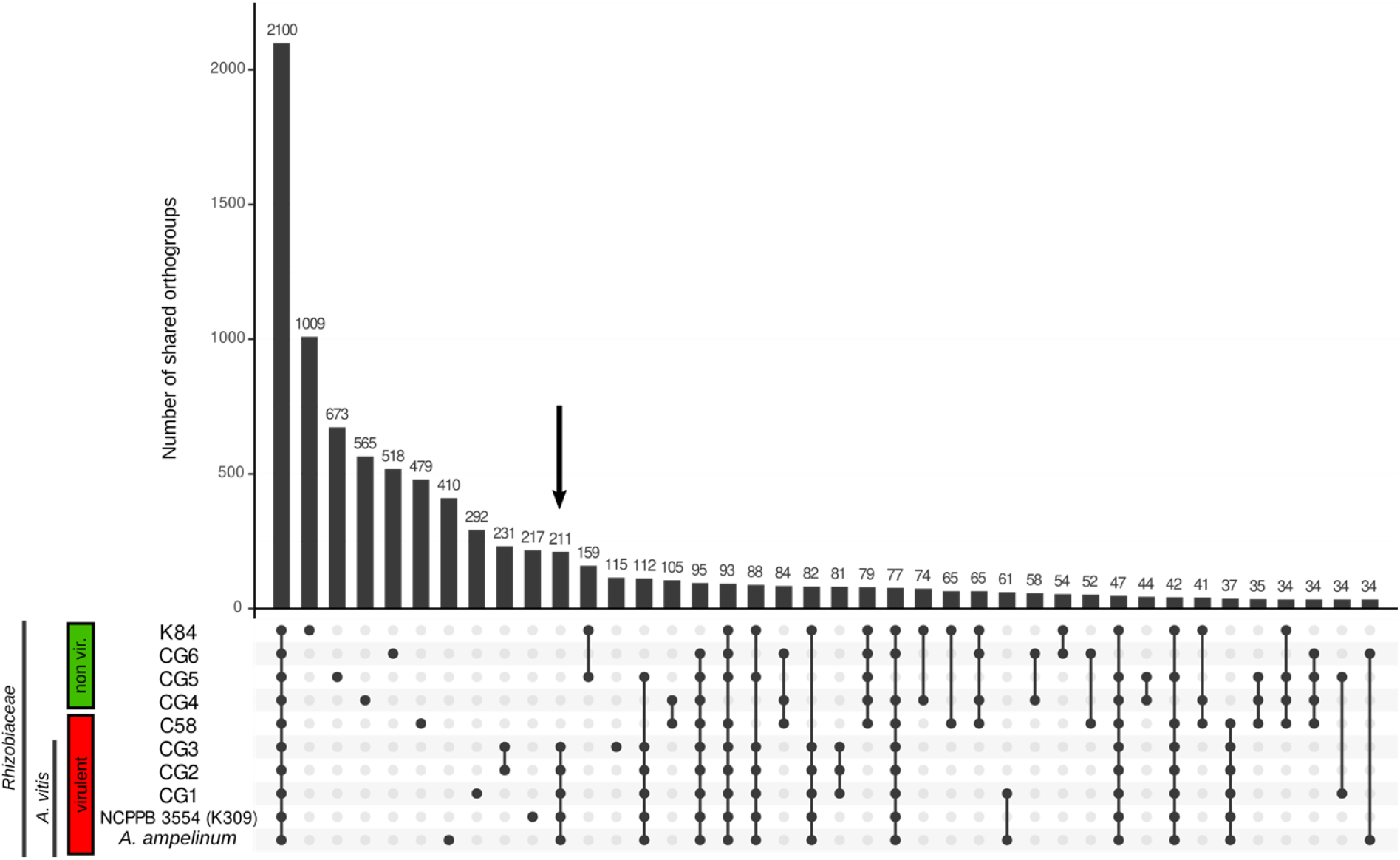
Shared and unique orthologous groups encoded by the sequences of the *Rhizobiaceae* genomes (CG1-CG6) and the references *A. ampelinum*, *A. vitis* NCPPB3554 (K309), *A. fabrum* C58, and *A. rhizogenes* K84. Each column represents the number of orthologous genes only shared by the genomes (black dots below each bar). The red box indicates virulent bacteria and the green box non-virulent. Less than 34 shared protein families are not shown.

### Relationships between isolates and reference Ti plasmids

The relationship between Ti-plasmids was investigated by aligning the potential pTi sequences of our *de novo* sequenced strains with each other and with the reference sequences of pTiS4 (vitopine/heliopine-type), pTiAg57 (octopine/cucumopine-type), pTiC58 (nopaline-type), and pTiK309 (octopine-type). The potential Ti-plasmids of the sequenced isolates CG1, CG2, and CG3 are more homologous to each other than to the reference pTi sequences. Among the three, CG2 and CG3 show a higher degree of homology to each other than to CG1 (fig. 3*A*). The alignments of CG1-CG3 to the reference pTis revealed that large parts of the CG1 contigs have strong homology (more than 99% identity) to long regions of pTiAg57 and almost the complete pTiK309 sequence (fig. 3*B*, dark green connecting lines). In particular, the *vir*-regions (fig. 3*B-D*, green bars) of the *de novo* sequenced *Allorhizobium* isolates (CG1-CG3) are highly homologous (100%) to the *vir-*regions of pTiAg57 and pTiK309 but much less to pTiS4 (orange connecting lines) and pTiC58 (red connecting lines). Furthermore, other contigs of CG1-CG3 encoding opine synthesis (fig. 3 purple bars), opine utilization and transport (fig. 3, grey bars), as well as plasmid replication and transfer (fig. 3, yellow bars) match to a high degree to pTiAg57.

**Fig. 3.**
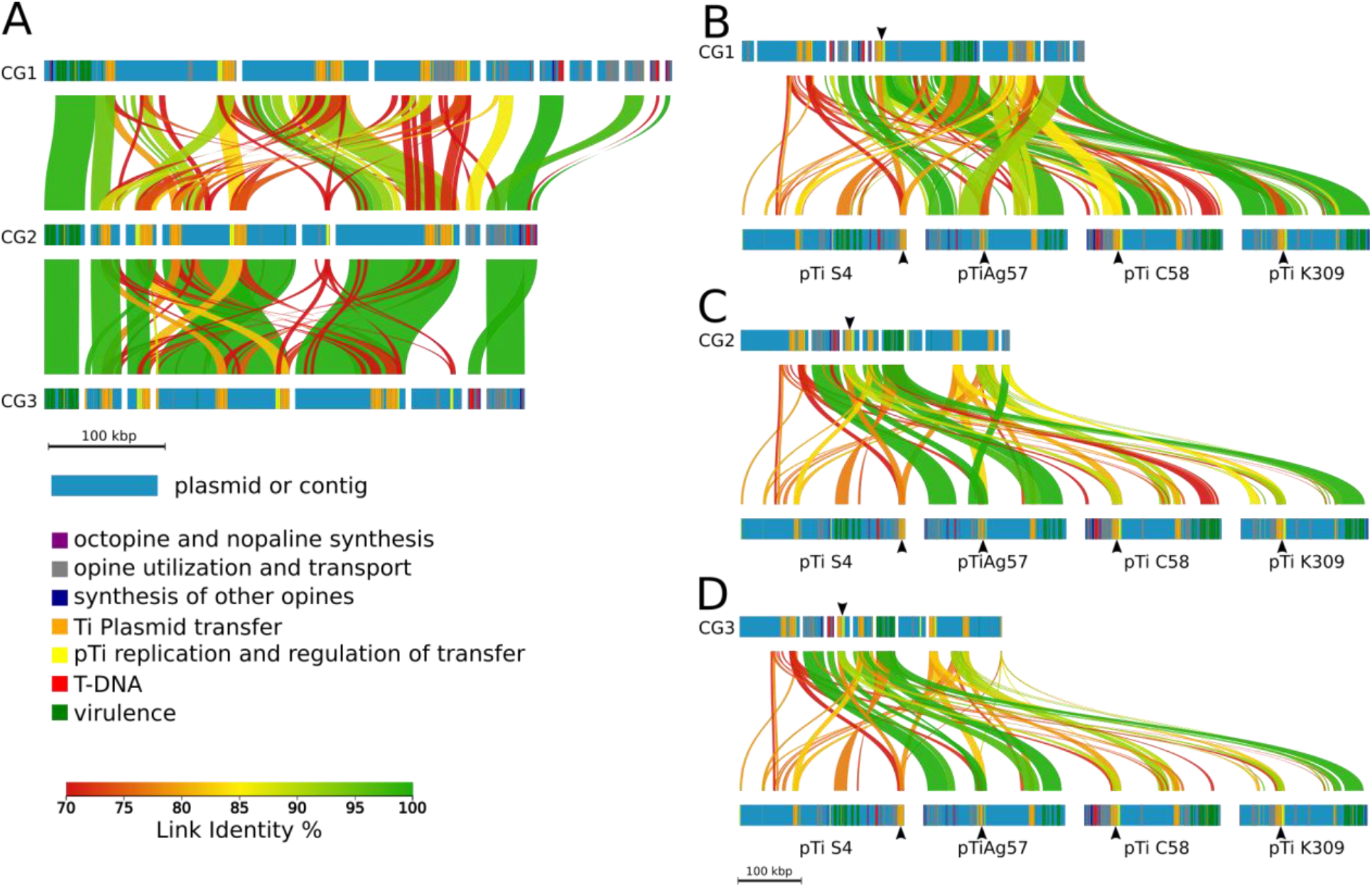
DNA alignments of Ti plasmid regions of the *de novo* sequenced isolates CG1-CG3 and three reference pTis. **(A)** Alignment of CG1, CG2, and CG3 and **(B-D)** of potential Ti-plasmids from CG1-CG3 to the reference Ti-plasmids of *A. ampelinum* (vitopine/heliopine-type), *A. fabrum* Ag57 (octopine/cucumopine-type), *A. fabrum* C58 (nopaline-type), and *A. vitis* NCPPB3554 (pTiK309) (octopine/cucumopine-type). Colored horizontal bars represent contigs of the potential Ti-plasmids and reference pTis. The position of predicted protein functions is marked by different colors. Homologous regions between the contigs are connected via vertical-colored lines and the color gradient shows % similarity (Link Identity %). Black arrow heads indicate the regions for Ti plasmid transfer (orange bars) and pTi plasmid replication (yellow bars) with highest homology to the reference plasmids.

### Prediction of Ti-Plasmid encoding protein families

To functionally describe the *de novo* sequenced genomes of the eight CG isolates, we compared the predicted protein sequences with those of the reference strains. The predicted proteins were clustered into orthologous groups (in short: orthogroups) by a set of amino acid sequences derived from a single common ancestor sequence for all the isolates (Emms and Kelly 2015). In total 21,431 unique predicted orthogroups were identified and for a selection of 15 plant associated bacteria, including the eight CG isolates and seven of the reference genomes, 495 orthogroups were shared by all the isolates (fig. 4). The genomes of the *Rhizobiaceae* had 349 additional unique orthogroups in common, and the taxonomically heterogeneous genomes of crown gall-associated bacteria shared only a single unique orthogroup. However, the subgroup of virulent bacterial genomes exclusively shared 28 orthogroups with each other.

**Fig. 4.**
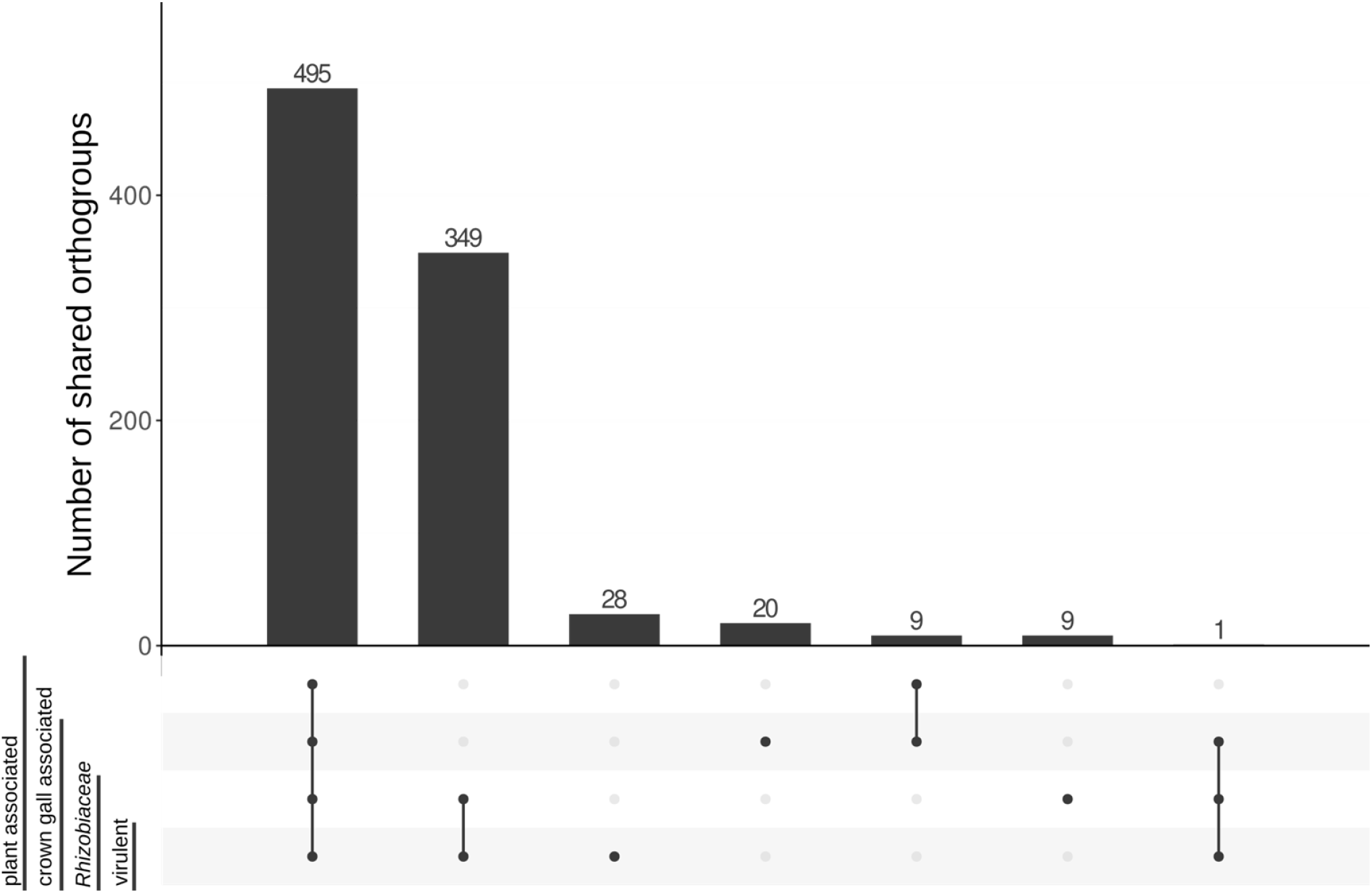
Shared and unique protein families among the selected genomes. These are grouped into i) plant associated, ii) crown gall associated, iii) Rhizobiaceae, and iv) virulent. The four groups consist of the following members i) *Pseudomonas cerasi*, *Rhanella aquatilis*, *Shpingomonas sp*. Leaf230, *Curtobacterium strain 6*, *Curtobacterium flaccumfaciens,* ii) *Pseudomonas* CG7 and *Rhanella* CG8, iii) non-virulent *Rhizobiaceae CG4-CG6*, and iv) *Agrobacterium fabrum C58, Allorhizobium ampelinum,* the virulent *Allorhizobium vitis* isolates C*G1-CG3*. A protein family is part of a group if it occurs in all members (black dots). A protein family is not part of a group if it occurs in none of the members. Protein families that occur in some representatives of a group are discarded.

The proteins known to be encoded by the Ti-plasmids (pTiC58, pTiS4, pTiK309, pTiAg57) of the reference bacteria *A. tumefaciens* C58, *A. ampelinum, A. vitis* NCPPB3554, and *A. fabrum* Ag57, respectively are listed in table 2 and assigned to all of our isolates CG1-CG8. The predicted protein families functioning in virulence and CG induction were present in the genomes of the three virulent isolates CG1-CG3 but largely absent in the genomes of the non-virulent bacteria (table 2A). An exception was VirG which is part of the sensory response system and therefore found in all analysed genomes in substantial numbers. Proteins related to plasmid replication and transfer (Tra, Trb, RepABC) were not only present in the virulent CG1-CG3 but also in the non-virulent isolates CG4, CG5 and CG6, while they were missing in *Pseudomonas sp.* (CG7) and *Rhanella sp.* (CG8) (table 2B). Protein families predicted to be associated with opine utilization such as opine catabolism (OoxA/B, NoxA/B, VoxA/B) occurred in all analysed genomes (table 2C) and were not unique to the virulent bacteria in contrast to those for opine anabolism (Vis, Cus, Ocs, Acs, Acsx). Proteins functioning in opine transport (e.g. AccC-E and OccPch, NocP) were found in large numbers. Taken together, virulence, opine degradation, and T-DNA encoded protein families were predominantly found in the genomes of the virulent *A. vitis* isolates. Those predicted to transfer and replicate Ti-plasmids were present in all *Rhizobiaceae* isolates and proteins for opine transport and catabolism occurred in all listed plant associated bacteria.

**Table 2.**
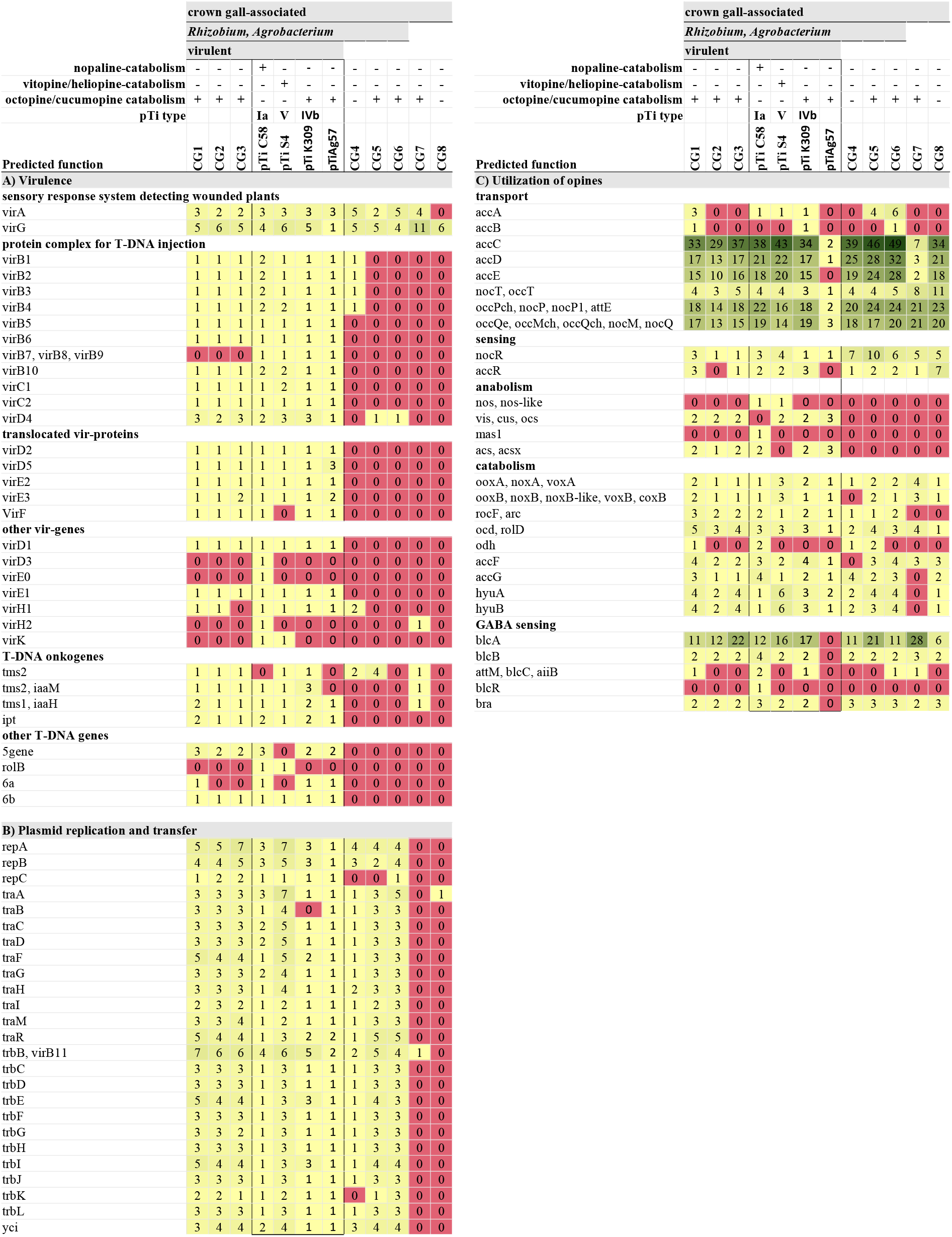
Number of predicted protein families encoded by the Ti plasmids of the reference bacterial strains *Agrobacterium fabrum* C58 (pTiC85) and LBA649 (pTiAg57), *Allorhizobium ampelinum* (pTiS4), *Allorhizobium vitis* NCPPB3554 (K309), and *A. fabrum* Ag57 (pTiAg57) as well as the de novo sequenced draft genomes of the crown gall associated isolates CG1-CG8. GABA sensing genes are encoded on pAtC58 (NC_003064). Representative names of the predicted protein families are listed in the first column. Gene counts are from whole genome, not only from plasmids, except for pTiAg57 where only the plasmid sequence is known.

### Genetic basis for opine utilization match *in vitro* opine bioassay

All CG associated isolates, except CG4, contained representatives of the well-described Nox/Oox/Vox protein families, essential for opine degradation. Figure 5 visualizes the order and orientation of the predicted proteins in the vicinity of the subunits A and B ofOox/Nox/Vox (green and turquoise arrows) in the *de novo* sequenced genomes and the reference *A. vitis* NCPPB3554 (K309). The fragments of the virulent *A. vitis* isolates CG1-CG3 showed exactly the same structure as the reference. This included the NoxA/B, OoxA/B, and VoxA/B protein families (fig. 5, green and turquoise arrows), other protein families related to opine catabolism (fig. 5, pale green arrows) and those for opine transport (fig. 5, blue arrows) upstream of OoxB/NoxB/VoxB (fig. 5, green arrows). Isolate CG1 harboured a second cluster with additional genes between the transport protein family and OoxB/NoxB/VoxB. Growth assays with the virulent CG1-CG3 isolates in liquid AB salt medium supplemented with opines as sole carbon and nitrogen source, confirmed the function of the opine catabolism clusters (supplementary table S3) The isolates utilized octopine better than nopaline pointing to the presence of octopine catabolism gene clusters in their genomes.

**Fig. 5.**
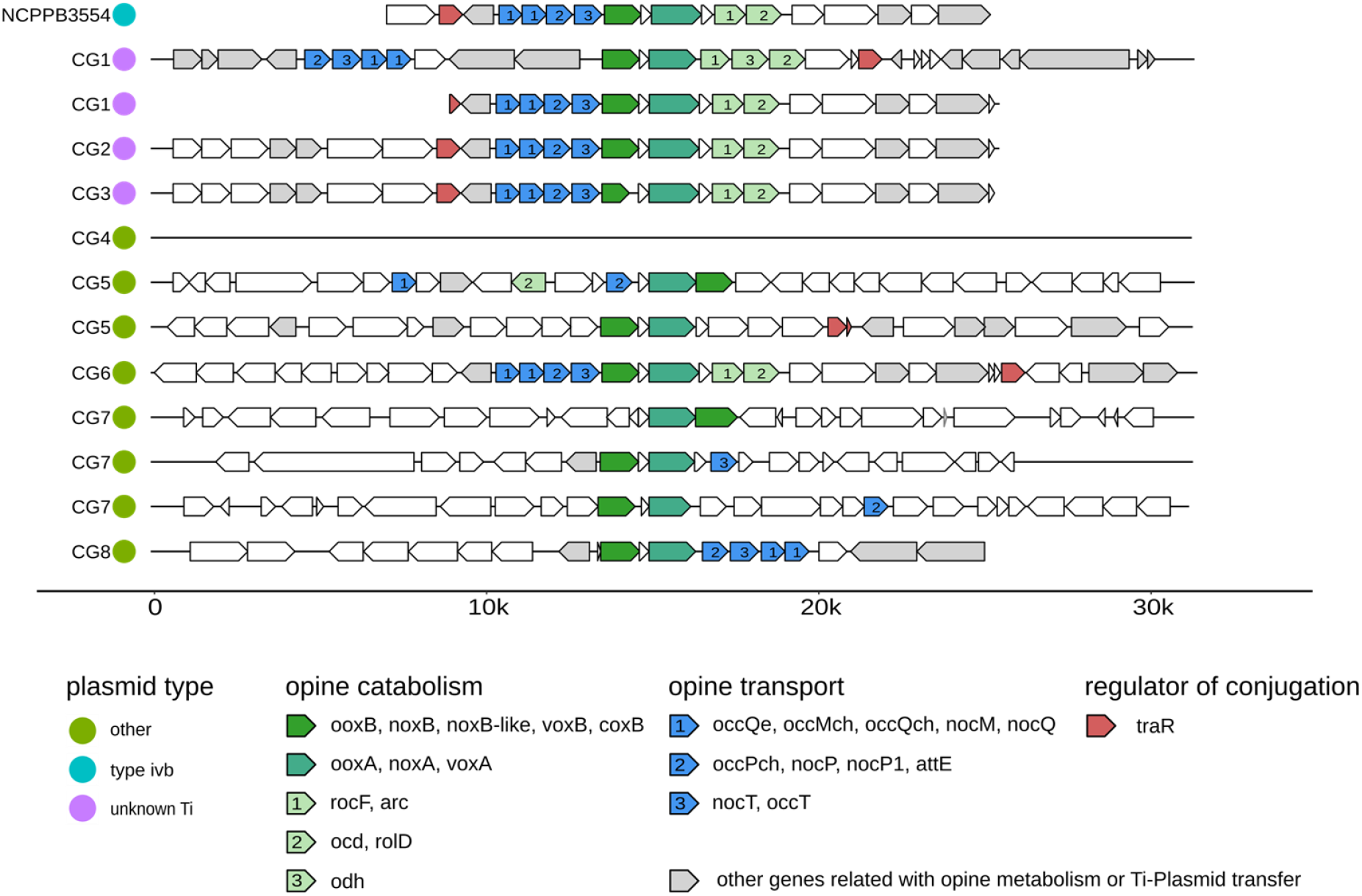
Regions in the *de novo* sequenced genomes of the CG1-CG8 isolates from grapevine crown galls that contain orthologs to both ooxA and ooxB, compared to the octopine catabolism cluster of pAtK309. The horizontal black lines represent a part of a contig centered around the ooxA/B, noxA/B, voxA/B genes. Colored arrows symbolize the orientation of coding regions including known annotations of PROKKA. Green colors symbolize opine catabolism; blue, opine transport; red, regulator of plasmid conjugation, and grey, other genes related with opine metabolism or Ti-plasmid transfer. The number within the arrows specify the gene function within a cluster.

The three non-virulent *Rhizobiaceae* isolates CG4, CG5, and CG6 showed a different behaviour concerning opine utilization. *A. divergens* (CG4) did not grow, but *Rhizobiaceae sp. (*CG5) grew well and *A. rosae* (CG6) weakly in liquid AB salt medium supplemented with octopine (supplementary table S3). Accordingly, the opine degradation region of CG4 lacked all substantial components (Fig. 5, black line) while the two clusters of CG5 harboured the essential genes, however, in a different order as compared to the virulent isolates CG1-CG3, except of the Nox/Oox/Vox proteins (fig. 5). CG6 possessed the same opine cluster structure as the *A. vitis* NCPPB3554 (K309) reference strain and the virulent isolates *A. vitis* CG1-CG3 but grew only weekly in opine containing liquid medium (supplementary table S3). A closer inspection of CG6 revealed additional predicted protein sequences that were homologous to those of the isolate CG1 and the reference plasmid pTiC58 of *A. fabrum* C58 (fig. 6A). In the reference plasmid, the homologs were related to replication and regulation of transfer (yellow bars), Ti plasmid transfer (orange bars), and opine utilization and transport (grey bars). Regions related to virulence (green bars) and the T-DNA (red bars) were not detected in the isolate CG6 genome but in the isolate CG1. This underlined the finding that CG6 was unable to induce CGs (supplementary fig. 1) but able to metabolize octopine, although not very well (supplementary table S3). Therefore, we compared the opine utilization and transport regions of CG6 also with the non-virulent agrocinopine/nopaline-type plasmid pAtK84b of *A. rhizogenes K84* and the octopine/cucumopine-catabolic *A. fabrum* plasmid pAtAg67 of the narrow host range *A. fabrum* strain Ag57. We found higher sequence identity between those than between CG6 and the Ti plasmid of C58 (fig. 6B, green connecting lines between grey bars). Particularly, one contig of CG6 matched the cucumopine-catabolic region of pAtAg67 with 99.5% identity. Thus, it is likely that the opine utilization and transport region of CG6 belongs to an opine-catabolic plasmid, most likely a cucumopine-type, rather than to a Ti plasmid.

**Fig. 6.**
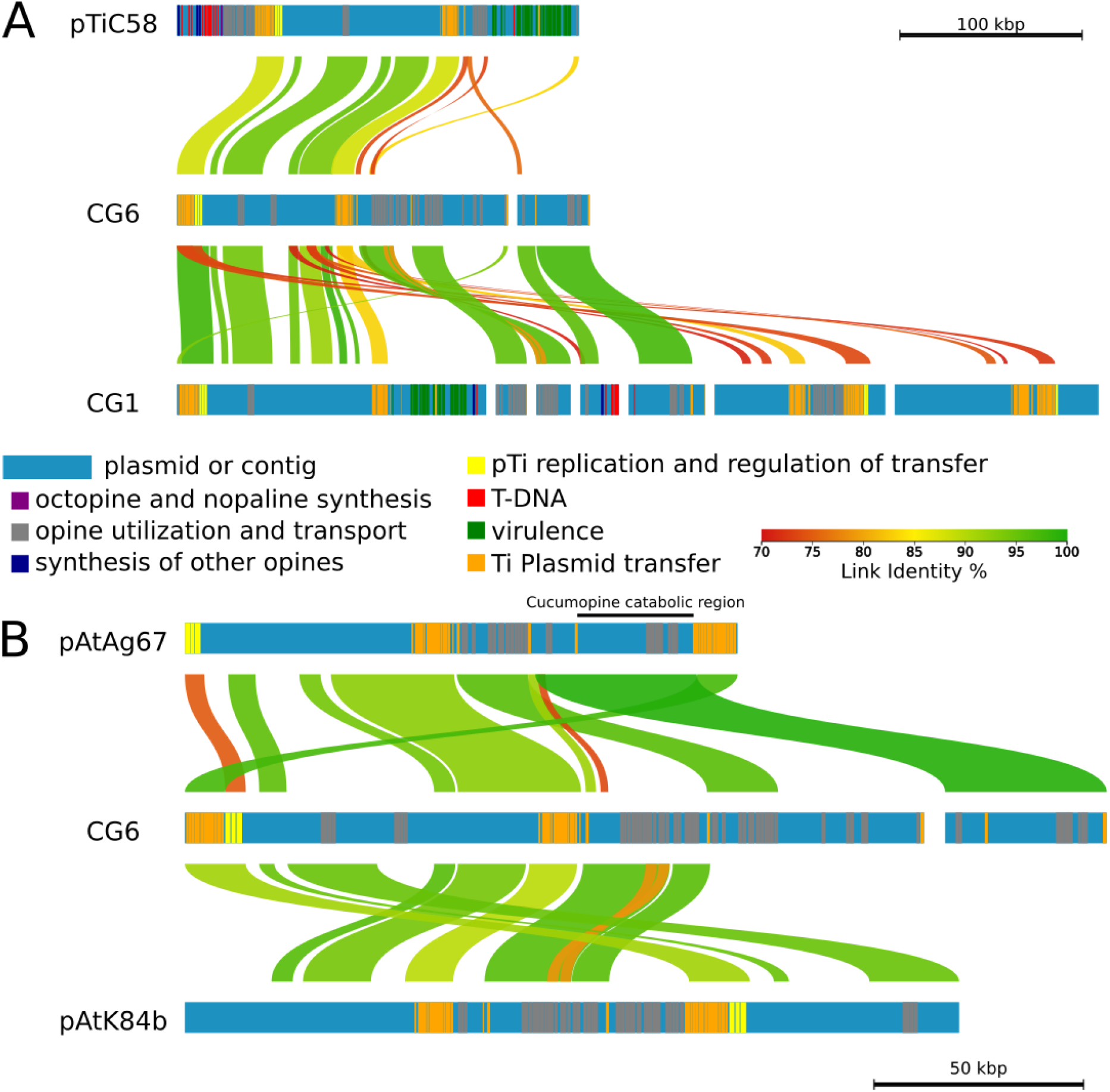
DNA alignment of putative plasmid sequences with reference plasmids. Colored horizontal bars represent the contigs and encoded protein families colored according to their function. Homologous regions between the sequences are connected via lines in a color gradient corresponding to % similarity (Link Identity %). **(A)** alignment of pTi regions from the reference strain *A. fabrum C58* and putative plasmid sequences of two isolates, the virulent *A. vitis* CG1 and non-virulent *A. rosae* CG6. **(B)** alignment of the opine-catabolic plasmid regions from the virulent reference *A. fabrum* Ag57 (pAtAg67), the non-virulent agrocinopine/nopaline-type plasmid of *A. rhizogenes K84* (pAtK84b) and putative plasmid sequences of the non-virulent *A. rosae* CG6. The cucumopine catabolic region of pAtAg67, according to Hooykaas et al. (2022) is indicated with a black bar.

The draft genome of the *Pseudomonas* isolate CG7 contained three regions with homology to the OoxA/B, NoxA/B, VoxA/B protein families, none of them has all the genes like the reference (NCPPB3554; fig. 5). However, the CG7 isolate grew well in liquid AB salt medium containing octopine like *A. rosa* CG6 and despite the differences in the structure of the opine clusters to those of the virulent isolates CG1-CG3 (supplementary table S3). In the *Rahnella* CG8 genome the DNA sequences encoding the OoxB/NoxB/VoxB protein family were also present in the classical orientation upstream of OoxA/NoxA/VoxA (fig. 5). However, in contrast to the opine-utilization clusters of CG1-CG3, the opine-transporter genes (fig. 5, blue arrows) in CG8 were downstream of OoxA/B, NoxA/B, VoxA/B and the isolate was not able to metabolize opines (supplementary table S3). Taken together, our study reveals that the non-virulent CG-associated bacteria *A. rosae* CG6 and *Pseudomonas* CG7 possess functional opine transport and catabolism sequence regions.

## Discussion

Virulent *Allorhizobium vitis* strains generate a new ecological niche in plants by transferring a T-DNA into the plant genome that leads to neoplastic growth of grapevine tissue so-called crown galls (CGs). In contrast to normal stem tissues, CGs form a sink tissue characterized by a hypoxic environment and accumulation of sugars, amino acids, and opines (Deeken, et al. 2006) which are exclusively produced by T-DNA harbouring host cells. In addition to the pathogen, a seasonally stable microbiota resides in the nutrient-rich CG environment (Faist et al. 2016).

This study aims to unravel whether homologs of the protein-encoding genes typically located on the Ti-plasmid of virulent agrobacteria are also found in non-virulent bacterial members of the CG bacterial community. To identify these genes in CG associated bacteria, we included genomes of non-tumour-associated plant bacteria in our analysis. Thereby we focussed on the functions: virulence, plasmid replication and transfer (conjugation), as well as opine utilization summarized in table 2. The distribution of the underlying genes may allow us to draw conclusions regarding the importance of their role in CG ecology.

### Virulence is restricted to *A. vitis* in the CG community

Three out of the eight *de novo* sequenced bacterial isolates (CG1, CG2, CG3) analysed in this study belonged to the taxonomic group of *Allorhizobium* . They were isolated from different grapevine CGs and in an infection assay they induced neoplastic growth on *in vitro* cultivated grapevines. The sequences comprising the chromosomes of CG1-CG3 were largely homologous to those of the well characterized strain *A. ampelinum* (fig. 1). Concerning the Ti-plasmid sequences the three genomes showed a higher homology to each other than to the reference Ti-plasmids of *A. ampelinum* (vitopine/heliopine-type), *A. fabrum* Ag57 (octopine/cucumopine-type), *A. fabrum* C58 (nopaline-type), and *A. vitis* pTiK309 (octopine-type fig. 3). The Ti-plasmid pTiAg57 was more similar to the putative plasmid contigs of CG2 and CG3, than any of the other reference plasmids (pTiS4, pTiC58, pTiK309). These findings support the idea of i) an independent propagation of th chromosomes and Ti-plasmids, as previously suggested (Slater et al. 2009) and ii) a faster ecological specification of plasmids encoding virulence in contrast to chromosomes encoding mainly housekeeping functions (Weisberg et al. 2020). Both factors must be considered for the development and interpretation of diagnostic tests targeting the CG disease.

Essential for CG development are the virulence regions located on the Ti-plasmid, that are involved in the process of T-DNA transformation and have previously been summarized for *A. fabrum C58* (Pitzschke and Hirt 2010). Our *A. vitis* isolates CG1-CG3 shared similar predicted protein sequences associated with virulence of the Ti plasmids of *A. fabrum* C58 (pTiC58), *A. ampelinum* (pTiS4), *A. vitis* NCPPB3554 (pTiK309) and *A. fabrum* Ag57 (pTiAg57). Among those are the proteins known to be important for (i) recognition of the host cell and induction of Vir gene expression (VirA, VirG), (ii) the type IV secretion system (T4SS: VirB2, VirB5, VirB6, VirD4, etc.) for the transfer of the T-DNA (Tra and trb genes), and (iii) T-DNA integration (VirC2, VirE2, VirE3, VirD5, etc.) into the plant genome (table 2A).

The VirA and VirG protein families involved in host recognition were detected in all but CG8 (no VirA) *de novo* sequenced virulent and non-virulent genomes as well as the three reference genomes. For *A. fabrum* C58 it is known that the ubiquitously expressed two-way component phospho-relay system VirA/VirG detects and transduces signals of wounded plants and the phosphorylated VirG activates transcription of the Vir operon (Brencic and Winans 2005; Wise and Binns 2015). Nevertheless, VirG belongs to a large family of positive regulators responding to external phenolics, monosaccharides and pH (Winans et al. 1986), indicating that the predicted protein family plays multiple roles besides virulence regulation.

The agrobacterial VirB/VirD4 system forms the T4SS, dedicated to deliver the T-DNA and effector proteins into plant cells in the course of infection. Most protein homologs of the T4SS encoded on the three reference Ti plasmids (pTiC58, pTiS4, pTiK309,pTiAg57) were present in the virulent *A. vitis* isolates (CG1-CG3). However, our virulent isolates harbour most likely a T4SS variant distinct from the one of the reference Ti plasmids because sequences encoding VirB7, VirB8, VirB9 were not present in the genome data of CG1-CG3. These Vir proteins have stabilizing functions and are either located in or at the outer (VirB7, VirB9) as well as inner (VirB8) bacterial membrane (Gordon and Christie 2014).

The agrobacterial factors translocated into host cells by the T4SS are essential for T-DNA integration and include the VirD2-T-DNA complex, VirD5, VirE2, VirE3, and VirF (Vergunst et al 2005; Schrammeijer et al. 2003). These protein families have different functions and were present in the genomes of our virulent isolates CG1-CG3. Taken together, the genome sequences of our *A. vitis* isolates CG1-CG3 harbour the genetic repertoire required to cause CG disease (supplementary fig. 1).

### All *Rhizobiaceae* isolates harbour the machinery for Ti-plasmid conjugation

CGs are a perfect niche for plasmid conjugation within a bacterial population (Dessaux and Faure 2018). The proteins necessary for replication (RepABC) and transfer (Tra, Trb) of Ti-plasmids have been previously described in detail (Lang and Faure 2014). Representatives of these protein families existed in our virulent and non-virulent *Rhizobiaceae* genomes (table 2B), but not in the - isolates CG7 (*Pseudomonas sp.*) and CG8 (*Rahnella sp.*) that were also part of the CG microbiota (Faist et al. 2016). The genomes of the three virulent *A. vitis* isolates (CG1-CG3) each contained a region (fig. 3*B-D*, black arrow heads) coding for pTi replication (yellow bars) and transfer (orange bars) which had highest homology to pTiAg57 of *A. fabrum* Ag57 (≤ 100%, green connecting lines), less homology to pTiC58 of *A. fabrum* C58, pTi309 of *A. vitis* NCPPB3554 (≤ 90%, pale green connecting lines), and least to pTiS4 of *A. ampelinum* (≤ 80%, orange connecting lines). Among the non-virulent *Rhizobiaceae* isolates, CG4 (*A. divergens*) and CG5 (*Rhizobium sp.*) seem to lack the sequence encoding replicase RepC based on the orthologue prediction (table 2) . However, the automatic gene annotation by NCBI revealed multiple genes classified as repC. This indicates the limitation of our gene assignment method, which can lead to false negatives. The also non-virulent *Rhizobiaceae* isolate CG6 (*A. rosae*) contained an almost identical sequence region to the virulent isolate *A. vitis* CG1 (fig. 6A, green connecting line) involved in plasmid replication (yellow bars) and transfer (orange bars) suggesting the presence of a plasmid, but not of a virulent Ti-plasmid. Regions involved in virulence (green bars) and T-DNA transfer (red bars), found in the genomes of C58 and CG1 were lacking in CG6.

*Quorum sensing* and *quorum quenching* regulate conjugation of Ti-plasmids between bacteria and horizontal transfer of the T-DNA into the host genome (Dessaux and Faure 2018). Opines synthesized by T-DNA transformed tumour cells play a key role in bacterial conjugation since they activate the conjugal transfer (*tra*) genes (Mel and Mekalanos 1996). Sequences of the predicted protein families involved in *quorum sensing* (TraI, TraM TraR) were present in all virulent and non-virulent *Rhizobiaceae* genomes (table 2B), but not in *Pseudomonas* sp. (CG7) and *Rahnella sp.* (CG8). The fact that similar *quorum sensing* genes existed in the virulent and non-virulent *Rhizobiaceae* isolates raised the possibility of cross species communication. This seems likely between our virulent *A. vitis* (CG2) and non-virulent *Pseudomonas* (CG7) isolates as well as between *A. vitis* (CG3) and *Rahnella* (CG8) because these resided in the same tumour B and tumour C, respectively (supplementary table S3). Representatives of the lactonase protein families (AttM/BlcC, AiiB) which degrade the bacterial *quorum sensing* signal molecule N-acyl homoserine lactone and thus are functioning in *quorum quenching* were found in the virulent *A. vitis* (CG1), the non-virulent *A. rosae* (CG6), and *Pseudomonas* (CG7; table 2C). *Quorum quenching* reduces the Ti-plasmid transfer frequency and thereby the host transformation events by virulent agrobacteria (Haudecoeur et al. 2009; Haudecoeur and Faure, 2010; Lang et al. 2016). Since *A. vitis* CG1, *A. rosae* CG6 and *Pseudomonas* CG7 were from different CGs they cannot negatively influence plasmid conjugation among each other or between *Rahnella* (CG8) and *A. vitis* (CG3) and *Pseudomonas* (CG7) and *A. vitis* (CG2). Thus, the latter two isolate pairs of tumour B and C, each have the genetic repertoire to communicate via *quorum sensing* signals and transfer genetic material.

### Opine utilization plays a key role in CG colonization

The possibility for non-virulent bacteria to acquire the ability for opine utilization is advantageous for them in the CG environment. An overview of the genes involved in opine utilization by agrobacteria is provided in Vladimirov et al. 2015 and table 2C. In CGs, different opines can coexist (Petit et al. 1983) and their recognition is conferred by various periplasmic binding proteins (PBPs). The PBPs OccT and NocT can recognize nopaline and octopine, respectively or in the case of NocT even both opines (Vigouroux et al. 2017). Association of PBPs with ATP binding cassette transporters (OocQ, NocQ, OocM, NocM, OccP, NocP) enables the import of opines into bacterial cells (Zanker et al. 1992). In our study NocT, OccT homologs were present in all crown gall-associated genomes and representatives of the ABC transporters (OccPch, NocP1 and OccQe, OccMch, etc.) even in high copy numbers (table 2C). A similar ubiquitous occurrence showed protein families for opine sensing (NocR, AccR). These function in transcriptional activation of opine uptake and catabolism genes (Subramoni et al. 2014). Hence, all crown gall-associated bacteria harbour the genetic potential for opine uptake and sensing.

Ti plasmids carry the genes for opine synthesis (*nos*, *ocs*, *vis*, cus, acs, etc.) by plant cells as well as the corresponding catabolism genes (*noxA*/*noxB*, *ooxA*/*ooxB*, rocF etc.). The octopine synthase gene sequence (*ocs*) is similar to the one of vitopine synthase (*vis*) and therefore joins the same protein family while the nopaline synthase gene (*nos*) forms a separate family (Canaday et al. 1992). We found homologs of the opine synthases Ocs, Vis, Cus, and Acs/AcsX but not Nos, Nos-like, and Mas1 in the genomes of the virulent isolates GC1-GC3 only (table 2C). Since no Nos/Nos-like homologs were found in the genomes of isolates CG1-CG3 and these metabolized octopine much better than nopaline (supplementary table S3) we suggest that our three virulent *A. vitis* isolates harbour an octopine/vitopine-type Ti plasmid rather than a nopaline-type.

In contrast to anabolism, which was restricted to the virulent isolates, all analysed crown gall-associated bacterial genomes harboured sequences encoding enzymes for opine catabolism (table 2C). The protein families of OoxA/NoxA and OoxB/NoxB were found in all genomes, except of the isolate CG4 (*A. divergens*) which lacked OoxB/NoxB. OoxA/NoxA together with OoxB/NoxB are two soluble polypeptides, which are both required for the first step of opine utilization (Zanker, et al. 1994) and may explain why CG4 could not metabolise nopaline or octopine (supplementary table S3). Furthermore, the arrangement of the genes encoding the proteins for opine catabolism is essential for an effective function (Zanker, et al. 1994). The order and orientation of the genes involved in opine sensing (*nocR*), uptake (*occQe, occPch, nocP*) and catabolism (*ooxB*, *rocF*, *ocd*) in the vicinity of *ooxB/ooxA* of the type IVb reference Ti-plasmid of *A. vitis* NCPPB3554 (fig. 5) was the same in our virulent isolates (CG1-CG3). It differed in the genomes of the non-virulent isolates CG4, CG5, CG7, and CG8, but not CG6. *Rhizobiaceae sp.* CG5 metabolized octopine well and harboured the essential catabolism genes *ooxA* and *ooxB* in the correct order and orientation but the other genes in some distance. The gene order and orientation in the genome of *A. rosae* CG6 was the same as in the virulent isolates. This points to a different event of acquisition for the opine transport and catabolism genes and might have an impact on opine utilization which was less effective by CG6 compared to the virulent isolates CG1-CG3. The sequences for opine transport and catabolism of CG6 revealed a high degree of similarity to the pAtK84b plasmid of the strain *A. rhizogenes* K84 and an even higher to the pAtAg67 plasmid from *A. fabrum* Ag57 (fig. 6B, grey bars). *A. rhizogenes* K84 harbours a nopaline-catabolic plasmid (pAtK84b) and *A. fabrum* Ag57 an octopine/cucumopine-catabolic plasmid (pAtAg67; Hooykaas et al., 2022) which lack the Vir and T-DNA regions like the genome of CG6. *A. rhizogenes* K84 is frequently used as biocontrol agents to prevent CG development (Clare et al. 1990). Thus, OC plasmids have an ecological impact on controlling CG disease severity, possibly via advantages in the competition for opines with virulent strains and the secretion of antimicrobial substances (Platt et al. 2014).

The *Pseudomonas* isolate CG7 contained two regions which showed the essential succession of *ooxA/noxA* and *ooxB/noxB* but lacked sequences for *arc, odh*, and *ocd/rolD* (fig. 5). Nevertheless, the *in vitro* growth assay confirmed utilization of octopine by *Pseudomonas* CG7. Previously it was shown that in CGs of grapevines bacteria other than *A. vitis* can utilize opines, including *Pseudomonas* strains (Canfield and Moore 1991; Nautiyal and Dion 1990; Moore et al. 1997; Wetzel et al. 2014; Eng et al. 2015). However, the molecular mechanism behind it is yet unknown. The *Rahnella* isolate CG8 harboured the essential genes for opine catabolism but those for opine transport were located downstream of *ooxA/noxA* and *ooxB/noxB* (fig. 5). Therefore, one might speculate that the reversed order of the essential opine-catabolic and uptake genes may prevent opine utilization by Rahnella (CG8). The reference genome of *Rahnella aquatillis* HX2 (No. 34 in table 1) does also not harbour the essential genes for opine utilization and was proposed to function as biocontrol in confining CG disease (Chen et al. 2007). Consequently, the non-virulent CG isolates *Rhizobiaceae sp.* CG6, *Pseudomonas* CG7, and *Rahnella* CG8 have the potential to control CG disease.

In root nodules, acquisition events have previously been suggested between Alpha-, Beta- and Gammaproteobacteria (Shiraishi et al. 2010; Ryu et al. 2020; De Meyer et al. 2016). Thus, an exchange of gene sequences between our alphaproteobacterium *A. vitis* (CG1, CG2, CG3), betaproteobacterium *Rahnella* (CG8), and the gammaproteobacterium, *Pseudomonas* (CG7) seems possible in CGs since CG2 and CG7 as well as CG3 and CG8 resided in the same CG. This might equip the non-virulent members of the microbial CG community with the machinery to utilize opines. Moreover, the transfer of the opine utilization machinery to beneficial plant bacteria could stabilize their population in an opine enriched environment. Taken together, in CGs, virulent *Rhizobiaceae* provide an opine-rich niche for themselves and other opine catabolizing bacteria as well.

## Conclusion

On a genetic level, gene duplication, rearrangements and interspecies horizontal gene transfer may be important for the dissemination of opine utilization among the CG-associated bacterial community. Our results highlight the distribution of sequences encoding proteins for opine catabolism (utilization of the CG-specific nutrient), but not for virulence (induction of the CG disease) among the bacterial community of the CG. The extent of CG development correlates with the number of transformation events and plant vigour. Grapevines developing small CGs show no growth limitations (Ferreira et al. 1992; Schroth et al. 1988) and the grapevines of this study did also not display an obvious phenotype. Opines are unique compounds produced by CGs which can only be metabolized by bacteria expressing enzymes for opine uptake and degradation. In contrast, other plant nutrients must be shared by the whole CG community. Competition for opines between virulent *A. vitis* and non-virulent bacteria could balance the composition of the microbial community in a CG and thus, promote or limit the degree of the CG disease. Therefore, a better understanding of the factors balancing the composition of the microbial community and providing a bacterial community, which is dominated by beneficial microbes, might lead to novel disease management strategies.

## Material and Methods

### Isolation and cultivation of bacteria

In 2011, 2012 and 2013 we isolated bacteria from CG material sampled in vineyards of Franconia, Germany. The grapevine cultivars consisted of the scion Cabernet Dorsa, Scheurebe and Müller Thurgau grafted onto the rootstocks 5BB and SO4 (NCBI, BioProject: PRJNA624984). CG material was ground (2 min, 30 Hz) using a ball mill (Retsch, Hannover, Germany) and 300 mg CG powder was suspended in super purified water (RotisolV high-performance liquid chromatography [HPLC] gradient grade; Roth). After incubation for 2 h at 28°C, the supernatant was used for 10-fold serial dilutions. Agar plates containing yeast extract broth (YEB; 0.5% [wt/vol] tryptone, 0.5% [wt/vol] yeast extract, 0.5% [wt/vol] sucrose, 1.23% [wt/vol] MgSO4 [AppliChem, Darmstadt, Germany], 1.5% [wt/vol] Agar-Agar Kobe I [Carl-Roth, Karlsruhe, Germany]) supplemented with 213 µM cycloheximide (CHX; Sigma-Aldrich, St. Louis, MO, USA) were used for bacterial growth at 28°C. By growing colonies on rifampicin containing yeast extract agar (RIF-YEA, 10 µg/ml) plates, spontaneous rifampicin-resistant derivatives were selected for tracking them in their natural environment. Single colonies were sub-cultured at least five-times on YEA-CHX-RIF plates and used for *de novo* shotgun sequencing.

### Virulence and opine growth assays

Virulence assays were performed by inoculating bacterial suspensions into *Vitis vinifera* stems as described by Faist et al. 2016, stems of four weeks old *Arabidopsis thaliana* (accession Col-0), and *Nicotiana benthamiana* plants according to (Gohlke et al. 2013). Pictures were taken using a CCD camera (Leica DFC500, Leica Microsystems GmbH) attached to a stereo microscope (Leica MZFLIII, Leica Microsystems GmbH). Opine utilization assays were performed in liquid AB minimal medium (K2HPO4 3 g/L; NaH2PO4 1 g/L; MgSO4-7H2O 0.3 g/L; KCl 0.15 g/L; CaCl2 0.01 g/L; FeS04-7H2O 2.5 mg/L; pH 7) supplemented with 1 mg/ml octopine or nopaline (Vigouroux et al. 2017) or with sucrose+NH_4_ and glycerol as controls. Bacterial growth was determined as optical density (OD_600_) and defined as follows: no growth, OD < 0.1; very weak growth, 0.1 ≤ OD < 0.2; weak growth 0.2 ≤ OD < 0.5; growth OD ≥ 0.5. The growth experiments with opines were repeated five times and the control experiments two (glycerol) to three (sucrose+NH_4_) times.

### Sequencing and identification of bacterial genomes

Eight bacterial CG isolates (CG1-CG8) were sequenced either using an Illumina Miseq (2014, CG1-CG2 and CG4-CG6, 2x250 bp V2 chemistry) or a NextSeq (2017, CG3 and CG7-CG8, 2x150bp mid-throughput v2 500/550 kit) after library preparation with the Nextera XT and 24 indices kits. Raw reads were quality filtered, corrected (Q30), and assembled using SPAdes 3.10.1 (Bankevich et al. 2012). The assembled sequences were screened for bacterial contaminations using blobtools (Laetsch and Blaxter 2017) with taxonomic assignment via BLAST (Altschul et al. 1990). Contigs are assembled DNA-sequences and in this study synonyms of scaffolds and nodes. On the contigs, rRNAs, tRNAs, genes (filtered open reading frames), and coding sequences (CDS) were annotated with PROKKA v1.12 (Seemann 2014) and Barrnap 0.9 (Torsten Seeman, https://github.com/tseemann/barrnap). Completeness of the genomes is indicated by the abundance of the 107 essential genes (Dupont et al. 2012). The overall genome coverage was calculated by mapping the original reads back onto the assembled contigs using Bowtie2 (Langmead and Salzberg 2012). The full length 16S rRNA gene sequences of the *de novo* draft genomes were taxonomically identified with the EZBioCloud database (Yoon et al. 2017). These 16S sequences were also matched with BLAST against the dataset of 16S v4 amplicon sequence variants (ASVs) by Faist et al. 2016 to account for their relative abundances in the whole bacterial community. We calculated a phylogenetic tree using BcgTree (Ankenbrand and Keller 2016), including our *Rhizobiaceae* isolates as well as 94 randomly selected *Rhizobiaceae* bacteria and four *Bradyrhizobium* genomes as an outgroup from EZBioCloud. The accession numbers for the raw reads at NCBI are JABAED000000000, JABAEE000000000, JABAEF000000000, JABAEG000000000, JABAEH000000000, JABAEI000000000, JABAEJ000000000, JABAED000000000 and JABAIN000000000 while for the annotated assemblies it is DOI: 10.5281/zenodo.3752520.

### Comparative genomics

A total of 34 reference genomes were used in addition to the eight genomes from this study. The references include 14 strains of *A. vitis*, two strains of *A. fabrum/tumefaciens*, eight strains of *A. rhizogenes*, six strains from the genus *Pseudomonas*, one from *Rahnella,* one from *Shpingomonas,* and two *Curtobacterium* strains. Strain details and accession numbers are listed in table 1 and supplementary table 1. Most of the *A. tumefaciens*, *A. vitis*, and *A. rhizogenes* references are described in (Weisberg et al. 2020) including a classification of their oncogenic plasmid and opine metabolism capabilities. Orthologous groups (OG) based on amino acid sequences of our isolates (CG1-CG8) and the reference genomes were identified by OrthoFinder (Emms and Kelly 2015). Gene names were transferred from the reference genomes to all genes of the cluster. Protein families including known Ti-plasmid proteins from the references were sorted into the following functional groups: A) virulence, B) plasmid replication and transfer, and C) utilization of opines (table 2). DNA sequences were aligned with lastZ (Harris 2007) and visualized with AliTV (Ankenbrand et al. 2017). For this comparison, two additional, recently described, plasmids were included, namely pTiAg57 and pAtAg67 (Hooykaas et al. 2022). Genes of these two plasmids were assigned to orthogroups by best BLAST hit (Altschul et al., 1990) with e-value below 10^-12^. For an unsupervised approach, shared and unique OGs of different bacterial species were displayed in UpSetR (Conway et al. 2017). For identification of putative opine clusters, we analysed the genomic neighbourhood of regions, where homologs of both OoxA/NoxA/VoxA and OoxB/NoxB/VoxB genes are found. We selected fragments of 15 kbp downstream and upstream of these genes for direct comparisons. If not indicated otherwise, gene functional descriptions are from the STRING database (https://string-db.org/, March 2019)

## Data Availability

The genome sequences obtained in this study have been deposited at NCBI with BioProject accession number PRJNA624984, annotated assemblies are available from Zenodo with doi:10.5281/zenodo.3752520.

## Acknowledgements

Special thanks go to Peter Schwappach (Bavarian Regional Office for Viticulture and Horticulture, Veitshoechheim, Germany) for providing grapevine plants and to Ernö Szegedi (Research Institute for Viticulture and Enology, Kecskemét, Hungary) for providing us with the chemical compounds octopine and nopaline. Many thanks go also to our colleagues from the University of Wuerzburg (in particular Gudrun Grimmer for library preparation for genome sequencing) and the Graduate School of Life Sciences from the University of Wuerzburg. We also want to thank Lisa Walther for her work and the data she contributed to this study and acknowledge Anne Müller and Lorenz Hoffmann, who help us to isolate bacteria that were sequenced in this study during their Master thesis (2012) and Diploma thesis (2013) at the University of Wuerzburg, respectively. Finally, we thank Rainer Hedrich (University of Wuerzburg, Germany) for financial support during this study.

## Supplementary Material

**Supplementary Fig. 1.**
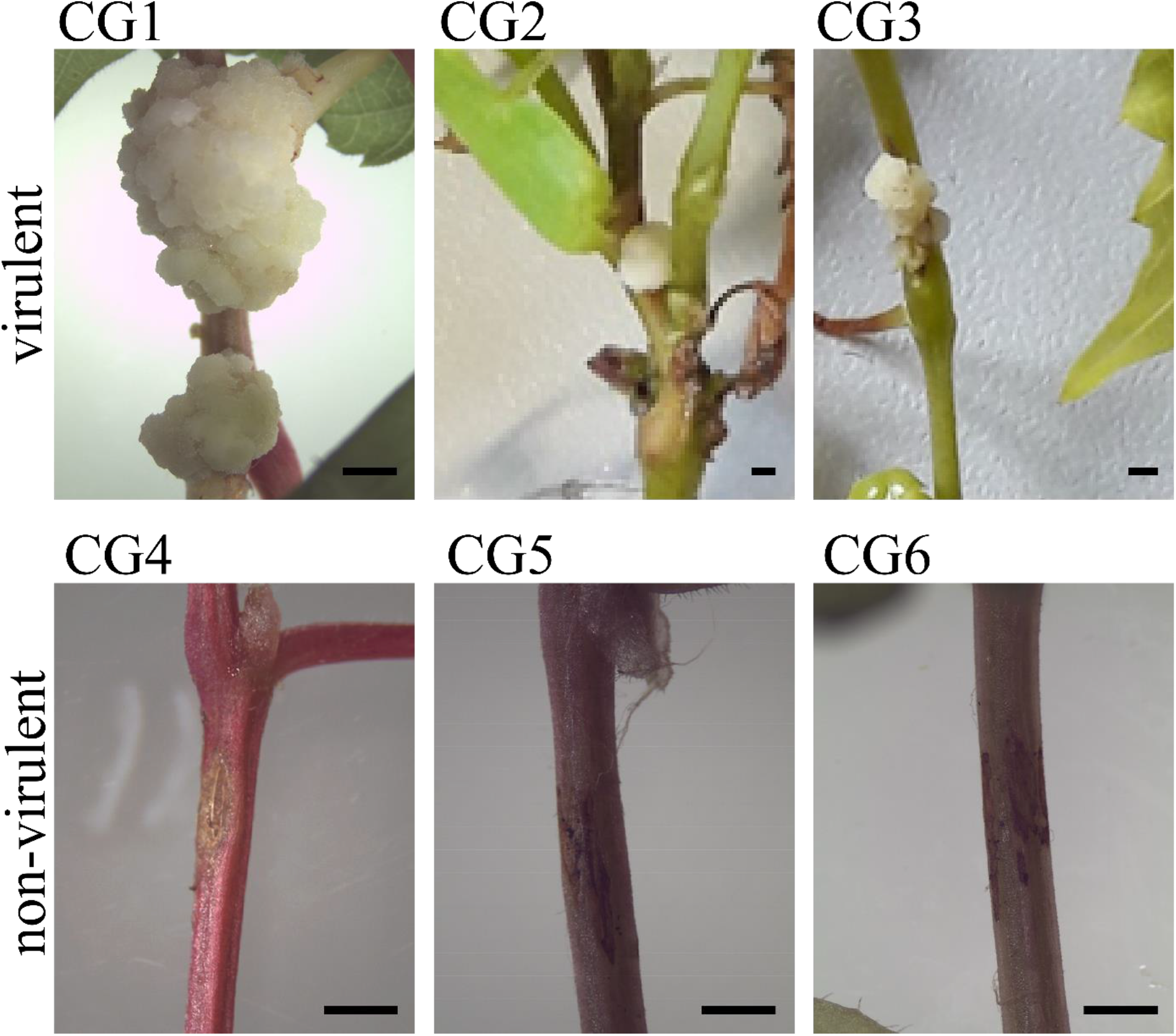
Virulence assay with the bacterial isolates CG1-CG6 and grapevine seedlings. Upper panel: Inoculation of the virulent isolates (CG1-CG3) caused crown gall formation in grapevine stems. Lower panel: Inoculation of the non-virulent isolates (CG4-CG6) induced no crown gall development at the wounded areas of grapevine stems (arrows). Bars represent 0.5 cm.

**Supplementary Fig. 2.**
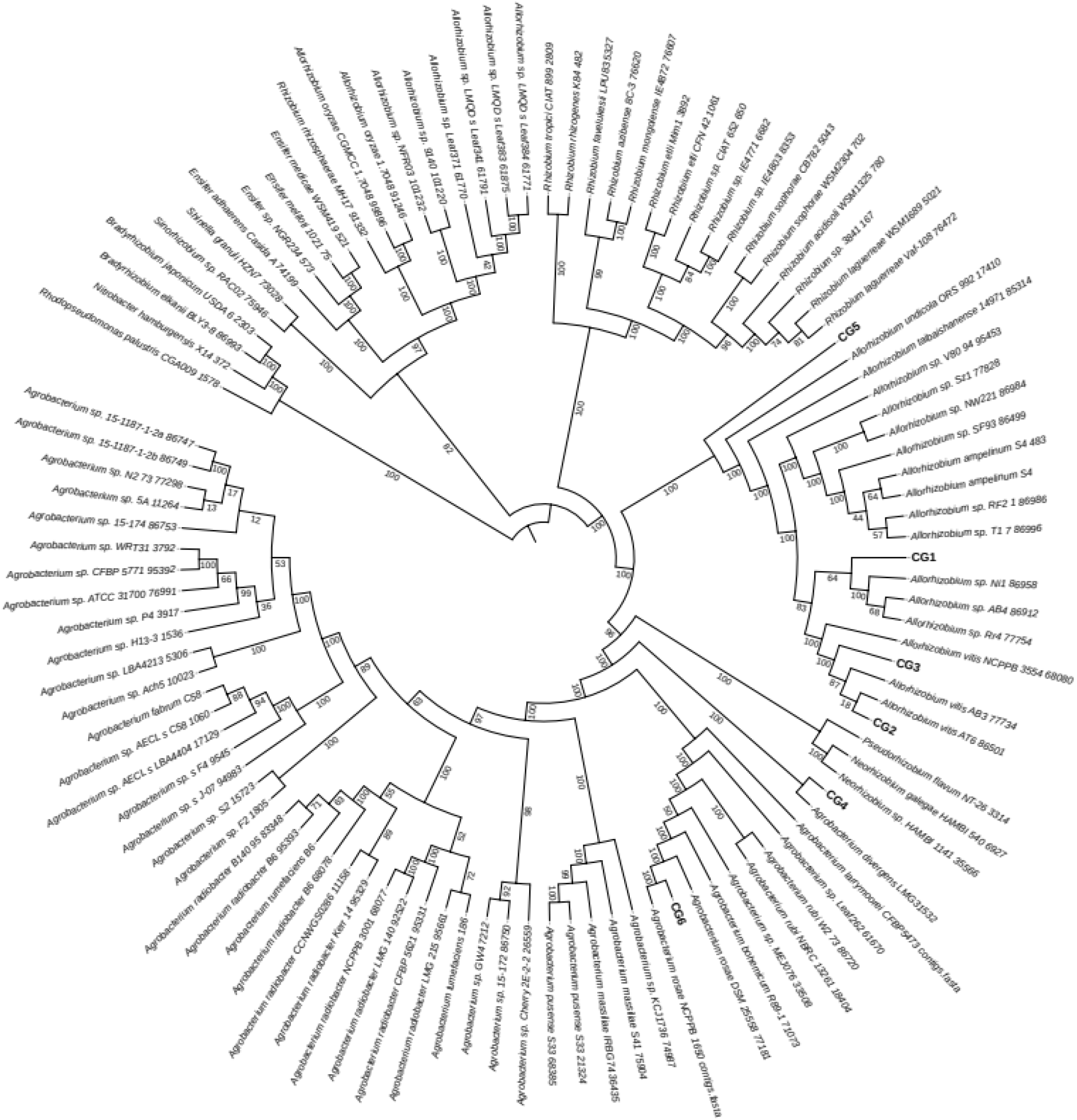
Phylogenetic tree based on 107 single copy housekeeping genes of members of the Rhizobiaceae family. The *de novo* sequenced isolates CG1-CG6 are integrated into a phylogenetic tree of Rhizobiaceae reference genomes from the EZBioCloud database.

**Supplementary Table S1.** Information on bacterial isolates of grapevine crown galls sampled from different vineyards in the Franconian region, Germany. Listed are 1. the characteristics of the isolates and 2. the relative abundance of 16S rRNA V4 amplicons in crown galls (% CG) and non-galled graft unions (% NG; Faist, et al 2017). Significant differences (p-value ≤ 0.05) between the graft unions with and without a crown gall were calculated according to one-way ANOVA followed by post hoc Tukey analysis for multiple testing.

| 1. Characteristics |  |  |  |  | 2. 16S rRNA V4 |  |  |
| --- | --- | --- | --- | --- | --- | --- | --- |
| Isolate | Tumor ID | Sampling date, vineyard | Reference strain | % | CG % | NG % | P-value |
| CG1, *Allorhizobium vitis* | A | Jan-09-2011, Himmelstadt, Germany | *Allorhizobium vitis*, NCPPB3554(T) | 99.9 | 11 | 0.3 | <0.0001 |
| CG2, *Allorhizobium vitis* | B | Jul-17-2013, Ravensburg, Germany | *Allorhizobium vitis*, NCPPB3554(T) | 99.9 | 11 | 0.3 | <0.0001 |
| CG3, *Allorhizobium vitis* | C | Oct-30-2013, Himmelstadt, Germany | *Allorhizobium vitis*, NCPPB3554(T) | 99.9 | 11 | 0.3 | <0.0001 |
| CG4, *Agrobacterium divergens* | B | July-17-2013, Ravensburg, Germany | *Rhizobium* sp., H13-3 | 98.8 | ND | ND | ND |
| CG5, *Rhizobiacea* sp. | D | Oct-24-2012, Sommerhäuser Reifenstein, Germany | *Rhizobium* sp., H13-3 | 98.6 | 0.8 | 0.8 | 0.9 |
| CG6, *Agrobacterium rosae* | E | Oct-30-2013, Himmelstadt, Germany | *Rhizobium* sp., Ch11(T) | 99.7 | 1.5 | 4.5 | 0.2 |
| CG7, *Pseudomonas* | B | July-17-2013, Ravensburg, Germany | *Pseudomonas* sp., DSM 13194(T) | 99.9 | 15.6 | 0.5 | <0.0001 |
| CG8, *Rahnella* | C | Oct-30-2013, Himmelstadt, Germany | *Rahnella* sp., Y9602 | 99.9 | 5.5 | 0.4 | <0.1 |

**Supplementary Table S2.** Features of the assembled draft bacterial genomes (CG1-CG8). N50 and N90 indexes list the length of the smallest contig that build 50% and 90%, respectively of the draft genomes. cont, contig.

| Draft genome | Total cont | Cont >1kb | 20 longest cont [Mb] | Total size [Mb] | N50 [kb] | N90 [kb] | GC [%] | Largest Kmer-coverage | Overall alignment rate [%] |
| --- | --- | --- | --- | --- | --- | --- | --- | --- | --- |
| *A. vitis* CG1 | 427 | 58 | 5.75 | 6.31 | 457 | 80.5 | 57.4 | 45.7 | 98.53 |
| *A. vitis* CG2 | 176 | 39 | 5.29 | 5.50 | 599 | 108 | 57.7 | 34.0 | 98.78 |
| *A. vitis* CG3 | 212 | 149 | 2.90 | 6.10 | 89.7 | 24.6 | 57.7 | 14.3 | 86.86 |
| *A. divergens* CG4 | 117 | 23 | 5.66 | 5.71 | 458 | 159 | 55.0 | 28.2 | 98.64 |
| *Rhizobiacea* sp. CG5 | 223 | 54 | 5.39 | 6.58 | 262 | 70.2 | 61.5 | 21.5 | 98.74 |
| *A. rosae*CG6 | 148 | 26 | 5.68 | 5.74 | 472 | 204 | 56.6 | 49.4 | 97.99 |
| *Pseudomonas* CG7 | 392 | 182 | 2.48 | 6.87 | 65.5 | 17.6 | 60.7 | 19.6 | 84.98 |
| *Rahnella* CG8 | 175 | 73 | 4.07 | 5.63 | 186 | 43.0 | 52.3 | 43.5 | 87.92 |

**Supplementary Table S3.** Bacterial growth assays in liquid medium with AB salts and supplemented with either nopaline, octopine, glycerol, or sucrose+NH_4_^+^ as sole C and N source. Optical density at 600 nm (OD_600_) was measured after 48 h. Mean values of OD_600_ represent 5 replicates of two experiments. As control served the *Agrobacterium* strains C58 utilising nopaline and B6 octopine. OD < 0.1, no growth; 0.1 ≤ OD < 0.2, very weak growth; 0.2 ≤ OD < 0.5, weak growth; OD ≥ 0.5, growth. Red and blue colours indicate presence of isolates in the same tumor.

| Tumor |  |  | Nopaline | Octopine | Glycerol | Sucrose+NH<sub>4</sub><sup>+</sup> |
| --- | --- | --- | --- | --- | --- | --- |
| **A** | *Allorhizobium vitis* | **CG1** | 0.37 +/- 0.12 | 0.58 +/- 0.16 | 1.42 +/- 0.17 | 0.85 +/- 0.00 |
| **B** |  | **CG2** | 0.14 +/- 0.02 | 0.87 +/- 0.11 | 1.97 +/- 0.06 | 3.40 +/- 0.42 |
| **C** |  | **CG3** | 0.26 +/- 0.07 | 1.14 +/- 0.42 | 1.98 +/- 0.03 | 2.10 +/- 0.07 |
| **B** | *Agrobacterium divergens* | **CG4** | 0.10 +/- 0.03 | 0.14 +/- 0.07 | 1.85 +/- 0.15 | 0.10 +/- 0.07 |
| **D** | *Rhizobiaceae sp.* | **CG5** | 0.15 +/- 0.03 | 1.01 +/- 0.25 | 1.92 +/- 0.04 | 2.68 +/- 0.04 |
| **E** | *Agrobacterium rosae* | **CG6** | 0.12 +/- 0.05 | 0.31 +/- 0.12 | 0.43 +/- 0.15 | 0.25 +/- 0.00 |
| **B** | *Pseudomonas* | **CG7** | 0.14 +/- 0.02 | 1.10 +/- 0.17 | 1.97 +/- 0.06 | 3.50 +/- 0.35 |
| **C** | *Rahnella* | **CG8** | 0.16 +/- 0.22 | 0.08 +/- 0.04 | 1.97 +/- 0.06 | 3.58 +/- 0.25 |
| - | *Agrobacterium tumefaciens* | **B6** | 0.14 +/- 0.05 | 0.71 +/- 0.16 | 1.56 +/- 0.02 | 2.45 +/- 0.14 |
| - |  | **C58** | 1.33 +/- 0.70 | 0.06 +/- 0.05 | 1.99 +/- 0.02 | 2.93 +/- 0.04 |
| - | non-inoculated | *ni* | 0.00 +/- 0.00 | 0.00 +/- 0.00 | 0.00 +/- 0.00 | 0.00 +/- 0.00 |

## Literature Cited

Altschul SF, Gish W, Miller W, Myers EW, Lipman DJ. 1990. Basic local alignment search tool. J Mol Biol 215: 403–410. doi: 10.1016/S0022-2836(05)80360-2

Ankenbrand MJ, Hohlfeld S, Hackl T, Förster F. 2017. AliTV—interactive visualization of whole genome comparisons. PeerJ Computer Science 3: e116. doi: 10.7717/peerj-cs.116

Ankenbrand MJ, Keller A. 2016. bcgTree: automatized phylogenetic tree building from bacterial core genomes. Genome 59: 783–791. doi: 10.1139/gen-2015-0175

Bankevich A, et al. 2012. SPAdes: a new genome assembly algorithm and its applications to single-cell sequencing. J Comput Biol 19: 455–477. doi: 10.1089/cmb.2012.0021

Bien E, Lorenz D, Eichhorn K. (1990) Isolation and characterization of *Agrobacterium tumefaciens* from the German vineregion Rheinpfalz. J Plant Dis Protect 97: 313–322

Bergeron J, Macleod RA, Dion P. 1990. Specificity of octopine uptake by *Rhizobium* and *Pseudomonas* strains. Appl Environ Microbiol. 56(5): 1453–1458. doi: 10.1128/aem.56.5.1453-1458.1990.

Brencic A, Winans SC. 2005. Detection of and response to signals involved in host-microbe interactions by plant-associated bacteria. Microbiol Mol Biol Rev 69: 155–194. doi: 10.1128/MMBR.69.1.155-194.2005

Burr TJ, Katz BH. 1983. Isolation of *Agrobacterium tumefaciens* biovar 3 from grapevine galls and sap, and from vineyard soil. Phytopathol 73: 163–165.

Canaday J, Gérad JC, Crouzet P, Otten L. 1992. Organization and functional analysis of three T-DNAs from the vitopine Ti plasmid pTiS4. Mol Gen Genet. 235(2-3):292–303. doi: 10.1007/BF00279373. PMID: 1465104.

Canfield ML, Moore LW. 1991. Isolation and characterization of opine utilizing strains of *Agrobacterium tumefaciens* and fluorescent strains of *Pseudomonas spp*. from rootstocks of Malus. Phytopathol 1(4): 440–443.

Chandrasekaran M, Lee JM, Ye BM, Jung SM, Kim J. et al. 2019. Isolation and Characterization of Avirulent and Virulent Strains of *Agrobacterium tumefaciens* from Rose Crown Gall in Selected Regions of South Korea. Plants (Basel) 25;8(11): 452. doi: 10.3390/plants8110452.

Chen J, Xie J. 2011. Role and regulation of bacterial LuxR-like regulators. J Cell Biochem 112: 2694–2702. Doi: 10.1002/jcb.23219

Chen F, Guo YB, Wang JH, Li JY, Wang HM. 2007. Biological Control of Grape Crown Gall by *Rahnella aquatilis* HX2. Plant Dis 91: 957–963. doi: 10.1094/PDIS-91-8-0957

Chilton WS, Petit A, Chilton M-D, Dessaux Y. 2001. Structure and characterization of the crown gall opines heliopine, vitopine and ridéopine. Phytochemistry 58 (1): 137–142, doi: 10.1016/S0031-9422(01)00166-2

Clare BG, Kerr A, Jones DA. 1990. Characteristics of the nopaline catabolic plasmid in *Agrobacterium* strains K84 and K1026 used for biological control of crown gall disease. Plasmid 23: 126–137. doi: 10.1016/0147-619x(90)90031-7

Conway JR, Lex A, Gehlenborg N. 2017. UpSetR: an R package for the visualization of intersecting sets and their properties. Bioinformatics 33: 2938–2940. doi: 10.1093/bioinformatics/btx364

De Meyer SE, et al. 2016. Symbiotic *Burkholderia* Species Show Diverse Arrangements of nif/fix and nod Genes and Lack Typical High-Affinity Cytochrome cbb3 Oxidase Genes. Mol Plant Microbe Interact 29: 609–619. doi: 10.1094/MPMI-05-16-0091-R

Deeken R, et al. 2006. An integrated view of gene expression and solute profiles of Arabidopsis tumors: a genome-wide approach. Plant Cell 18: 3617–3634. doi: 10.1105/tpc.106.044743

Dessaux Y, Petit A, Farrand S, Murphy P 1998. in The Rhizobiaceae: Molecular Biology of Model Plant-Associated Bacteria. Springer, pp. 173–197.

Dessaux Y, Faure D. 2018. Quorum Sensing and Quorum Quenching in *Agrobacterium*: A "Go/No Go System"? Genes (Basel) 9. doi: 10.3390/genes9040210

Dupont CL, et al. 2012. Genomic insights to SAR86, an abundant and uncultivated marine bacterial lineage. ISME J 6: 1186–1199. doi: 10.1038/ismej.2011.189

Ellis JG, Kerr A, Petit A, Tempe J. 1982. Conjugal transfer of nopaline and agropine Ti-plasmids —The role of agrocinopines. Molecular and General Genetics MGG 186: 269–274. doi: 10.1007/bf00331861

Emms DM, Kelly S. 2015. OrthoFinder: solving fundamental biases in whole genome comparisons dramatically improves orthogroup inference accuracy. Genome Biol 16: 157. doi: 10.1186/s13059-015-0721-2

Eng WWH, Gan HM, Gan HY, Hudson AO, Savka MA. 2015. Whole-genome sequence and annotation of octopine-utilizing *Pseudomonas kilonensis* (previously *P. fluorescens*) strain 1855-344. Genome Announc 3(3): e00463–15. doi:10.1128/genomeA.00463-15.

Escobar MA, Civerolo EL, Summerfelt KR, Dandekar AM. 2001. RNAi-mediated oncogene silencing confers resistance to crown gall tumorigenesis. Proc Natl Acad Sci U S A 98: 13437–13442. doi: 10.1073/pnas.241276898

Faist H, Keller A, Hentschel U, Deeken R. 2016. Grapevine (Vitis vinifera) Crown Galls Host Distinct Microbiota. Appl Environ Microbiol 82: 5542–5552. doi: 10.1128/AEM.01131-16

Ferreira JHS, van Zyl FGH, Staphorst JL 1992. *Agrobacterium tumefaciens* biovar 3 Responsible for Reduction in Yield and Vigour of Muscat d’Alexandrie. South African J. Enol. Vitic. S AFR J ENOL VITIC 13: 78–80.

Gan HM, et al. 2019. Insight Into the Microbial Co-occurrence and Diversity of 73 Grapevine (Vitis vinifera) Crown Galls Collected Across the Northern Hemisphere. Front Microbiol 10: 1896. doi: 10.3389/fmicb.2019.01896

Gan HM, Savka MA. 2018. One More Decade of *Agrobacterium* Taxonomy. Curr Top Microbiol Immunol 418:1–14. doi: 10.1007/82_2018_81.

Gelvin SB. 2010. Plant proteins involved in *Agrobacterium*-mediated genetic transformation. Annu Rev Phytopathol 48: 45–68. doi: 10.1146/annurev-phyto-080508-081852

Gohlke J, et al. 2013. DNA methylation mediated control of gene expression is critical for development of crown gall tumors. PLoS Genet 9: e1003267. doi: 10.1371/journal.pgen.1003267

Gordon JE, Christie PJ. 2014. The *Agrobacterium* Ti plasmids. Microbiol Spectrum 2(6): PLAS-0010-2013. doi: 10.1128/microbiolspec.PLAS-0010-2013.

Harris RS, editor. 2007. Improved pairwise alignment of genomic DNA. The Pennsylvania State University.

Haudecoeur E, Faure D. 2010. A fine control of quorum-sensing communication in *Agrobacterium tumefaciens*. Commun Integr Biol 3(2): 84–88. doi: 10.4161/cib.3.2.10429.

Haudecoeur E, et al. 2009. Different regulation and roles of lactonases AiiB and AttM in *Agrobacterium tumefaciens* C58. Mol Plant Microbe Interact 22: 529–537. doi: 10.1094/MPMI-22-5-0529

Hooykaas M, et al. 2022. Characterization of the Agrobacterium octopine-cucumopine catabolic plasmid pAtAg67. Plasmid 121. doi: 10.1016/j.plasmid.2022.102629

Klee H, et al. 1984. Nucleotide sequence of the tms genes of the pTiA6NC octopine Ti plasmid: two gene products involved in plant tumorigenesis. Proc Natl Acad Sci U S A 81: 1728–1732. doi: 10.1073/pnas.81.6.1728

Laetsch D, Blaxter M. 2017. BlobTools: Interrogation of genome assemblies [version 1; peer review: 2 approved with reservations]. F1000Research 6. doi: 10.12688/f1000research.12232.1

Lang J, et al. 2017. Fitness costs restrict niche expansion by generalist niche-constructing pathogens. ISME J 11: 374–385. doi: 10.1038/ismej.2016.137

Lang J, Gonzalez-Mula A, Taconnat L, Clement G, Faure D. 2016. The plant GABA signaling downregulates horizontal transfer of the *Agrobacterium tumefaciens* virulence plasmid. New Phytol 210: 974–983. doi: 10.1111/nph.13813

Lang J, Faure D. 2014. Functions and regulation of quorum-sensing in *Agrobacterium tumefaciens*. Front Plant Sci 5: 14. doi: 10.3389/fpls.2014.00014

Langmead B, Salzberg SL. 2012. Fast gapped-read alignment with Bowtie 2. Nat Methods 9: 357–359. doi: 10.1038/nmeth.1923

Mel SF, Mekalanos JJ. 1996. Modulation of Horizontal Gene Transfer in Pathogenic Bacteria by In Vivo Signals. Cell 87: 795–798. doi: 10.1016/s0092-8674(00)81986-8

Moore LW, Chilton WS, Canfield ML. 1997. Diversity of opines and opine-catabolizing bacteria isolated from naturally occurring crown gall tumors. Appl Environ Microbiol 63: 201–207. doi: 10.1128/aem.63.1.201-207.1997

Nautiyal CS, Dion P. 1990. Characterization of the Opine-Utilizing Microflora Associated with Samples of Soil and Plants. Appl Environ Microbiol 56: 2576–2579. doi: 10.1128/aem.56.8.2576-2579.1990

Petit A, et al. 1983. Further extension of the opine concept: Plasmids in *Agrobacterium rhizogenes* cooperate for opine degradation. Molecular and General Genetics MGG 190: 204–214. doi: 10.1007/bf00330641

Pitzschke A, Hirt H. 2010. New insights into an old story: *Agrobacterium*-induced tumour formation in plants by plant transformation. EMBO J 29: 1021–1032. doi: 10.1038/emboj.2010.8

Platt TG, Morton ER, Barton IS, Bever JD, Fuqua C. 2014. Ecological dynamics and complex interactions of *Agrobacterium* megaplasmids. Frontiers Plant Sci 5: doi: 10.3389/fpls.2014.00635

Platt TG, Fuqua C, Bever JD. 2012. Resource and competitive dynamics shape the benefits of public goods cooperation in a plant pathogen. Evolution 66: 1953–1965. doi: 10.1111/j.1558-5646.2011.01571.x

Ryu MH, et al. 2020. Control of nitrogen fixation in bacteria that associate with cereals. Nat Microbiol 5: 314–330. doi: 10.1038/s41564-019-0631-2

Schrammeijer B, den Dulk-Ras A, Vergunst AC, Jurado Jacome E, Hooykaas PJ. 2003. Analysis of Vir protein translocation from *Agrobacterium tumefaciens* using Saccharomyces cerevisiae as a model: evidence for transport of a novel effector protein VirE3. Nucleic Acids Res 31: 860–868. doi: 10.1093/nar/gkg179

Schroth M, McCain A, Foott J, Huisman O. 1988. Reduction in yield and vigor of grapevine caused by crown gall disease. Plant Disease 72: 241–246. doi:10.1094/PD-72-0241

Seemann T. 2014. Prokka: rapid prokaryotic genome annotation. Bioinformatics 30: 2068–2069. doi: 10.1093/bioinformatics/btu153

Shiraishi A, Matsushita N, Hougetsu T. 2010. Nodulation in black locust by the Gammaproteobacteria *Pseudomonas sp*. and the Betaproteobacteria *Burkholderia sp*. Syst Appl Microbiol 33: 269–274. doi: 10.1016/j.syapm.2010.04.005

Slater SC, et al. 2009. Genome sequences of three agrobacterium biovars help elucidate the evolution of multichromosome genomes in bacteria. J Bacteriol. 191(8):2501–11. doi: 10.1128/JB.01779-08

Subramoni S, Nathoo N, Klimov E, Yuan ZC. 2014. *Agrobacterium tumefaciens* responses to plant-derived signaling molecules. Front Plant Sci 5: 322. doi: 10.3389/fpls.2014.00322

Szegedi E. 2003 Opines in naturally infected grapevine crown gall tumors. Vitis 42 (1): 39–41, doi: 10.5073/vitis.2003.42.39-41

Szegedi E, Czako M, Otten L. 1996. Further evidence that the vitopine-type pTi’s of *Agrobacterium vitis* represent a novel group of Ti plasmids. Mol Plant Microbe Interact 9: 139–143. doi: 10.1094/MPMI-9-0139

Szegedi E, Czakó M, Koncz CS. 1988. Opines in crown gall tumours induced by biotype 3 isolates of *Agrobacterium tumefaciens*. Physiol Mol Plant Pathol 3(5): 237–247, doi: 10.1016/S0885-5765(88)80020-1

Vergunst AC, et al. 2005. Positive charge is an important feature of the C-terminal transport signal of the VirB/D4-translocated proteins of *Agrobacterium*. Proc Natl Acad Sci USA 102: 832–837. doi: 10.1073/pnas.0406241102.

Vladimirov IA, Matveeva TV, Lutova LA. 2015. Opine biosynthesis and catabolism genes of *Agrobacterium* tumefaciens and *Agrobacterium rhizogenes*. Russ J Genet 51: 121–129. doi: 10.1134/S1022795415020167

Vigouroux A, et al. 2017. Structural basis for high specificity of octopine binding in the plant pathogen *Agrobacterium tumefaciens*. Sci Rep 7: 18033. doi: 10.1038/s41598-017-18243-8

Weisberg A, et al. 2020. Unexpected conservation and global transmission of agrobacterial virulence plasmids. Science. 368. eaba5256. 10.1126/science.aba5256. doi: 10.1126/science.aba5256.

Wetzel ME, Kim KS, Miller M, Olsen GJ, Farrand SK. 2014. Quorum-dependent mannopine-inducible conjugative transfer of an *Agrobacterium* opine-catabolic plasmid. J Bacteriol 196: 1031–1044. doi: 10.1128/JB.01365-13

Wikipedia contributors. 2023. Ti plasmid. In: Wikipedia, The Free Encyclopedia. Retrieved from https://en.wikipedia.org/w/index.php?title=Ti_plasmid&oldid=1144023521

Winans SC, Ebert PR, Stachel SE, Gordon MP, Nester EW. 1986. A gene essential for *Agrobacterium* virulence is homologous to a family of positive regulatory loci. Proc Natl Acad Sci U S A 83: 8278–8282. doi: 10.1073/pnas.83.21.8278

Wise AA, Binns AN. 2015. The Receiver of the *Agrobacterium tumefaciens* VirA Histidine Kinase Forms a Stable Interaction with VirG to Activate Virulence Gene Expression. Front Microbiol 6: 1546. doi: 10.3389/fmicb.2015.01546

Yoon SH, et al. 2017. Introducing EzBioCloud: a taxonomically united database of 16S rRNA gene sequences and whole-genome assemblies. Int J Syst Evol Microbiol 67: 1613–1617. doi: 10.1099/ijsem.0.001755

Zanker H, von Lintig J, Schröder J. 1992. Opine transport genes in the octopine (occ) and nopaline (noc) catabolic regions in Ti plasmids of *Agrobacterium tumefaciens*. J Bacteriol. 174(3):841–9. doi: 10.1128/jb.174.3.841-849.1992.

Zanker H, et al. 1994. Octopine and nopaline oxidases from Ti plasmids of *Agrobacterium tumefaciens*: molecular analysis, relationship, and functional characterization. J Bacteriol 176: 4511–4517. doi: 10.1128/jb.176.15.4511-4517.1994

